# An oscillating MinD protein determines the cellular positioning of the motility machinery in archaea

**DOI:** 10.1101/2020.04.03.021790

**Authors:** Phillip Nußbaum, Solenne Ithurbide, James C. Walsh, Megha Patro, Floriane Delpech, Marta Rodriguez-Franco, Paul M.G. Curmi, Iain G. Duggin, Tessa E.F. Quax, Sonja-Verena Albers

## Abstract

MinD proteins are well studied in rod-shaped bacteria such as *E. coli*, where they display self-organized pole-to-pole oscillations that are important for correct positioning of the Z-ring at mid-cell for cell division. Archaea also encode proteins belonging to the MinD family, but their functions are unknown. MinD homologous proteins were found to be widespread in Euryarchaeota and form a sister group to the bacterial MinD family, distinct from the ParA and other related ATPase families. We aimed to identify the function of four archaeal MinD proteins in the model archaeon *Haloferax volcanii*. Deletion of the *minD* genes did not cause cell division or size defects, and the Z-ring was still correctly positioned. Instead, one of the mutations (Δ*minD4*) reduced swimming motility, and hampered the correct formation of motility machinery at the cell poles. In Δ*minD4* cells, there is reduced formation of the motility structure and chemosensory arrays, which are essential for signal transduction. In bacteria, several members of the ParA family can position the motility structure and chemosensory arrays via binding to a landmark protein, and consequently these proteins do not oscillate along the cell axis. However, GFP-MinD4 displayed pole-to-pole oscillation and formed polar patches or foci in *H. volcanii*. The MinD4 membrane targeting sequence (MTS), homologous to the bacterial MinD MTS, was essential for the oscillation. Surprisingly, MinD4 ATPase domain point-mutations did not block oscillation, but they failed to form pole-patches. Thus, MinD4 from *H. volcanii* combines traits of different bacterial ParA/MinD proteins.

## Introduction

The position of macromolecular assemblies is regulated spatially and temporally within cells. Archaea and bacteria require active mechanisms to precisely position large protein complexes. Many bacterial cells are polarized and bacterial proteins with diverse functions display spatiotemporal dynamics, such as localization to mid-cell, to cell poles or along the long axis of rod-shaped cells^1^,^2^. Proteins from the ParA/MinD family are widespread in bacteria^3^ and responsible for the correct spatiotemporal placement of many different molecules and structures, including the chromosome, cytoplasmic protein clusters, pili, flagella and chemosensory machinery^4–9^.

ParA and MinD belong to the P-loop NTPase superfamily, which can form dimers^10–13^. A hallmark of these ATPases is a deviant Walker A motif (KGGXXGKT), where the amino terminal lysine is involved in dimerization by binding to the phosphate of the ATP of the other subunit^3^,^14^. Dimerization leads to increased affinity for the surface or protein partners to which the respective ParA/MinD proteins bind^12^, which is the cytoplasmic membrane in case of MinD^11^. ParA forms dimers that can bind nonspecific dsDNA in a nucleotide-dependent manner^15^, while ATP binding stimulates polymerization of the protein^13^. Generally, ParA and MinD have a partner protein that accelerates ATPase activity upon binding, such as ParB or MinE, respectively^16^,^17^. For the Min system, molecular interactions, ATPase activity, and diffusion rates can work together to establish protein concentration gradients on cellular surfaces, manifesting as a dynamic and self-organized pattern within cells^10^,^18^. For the Par system, molecular interactions and the ATPase activity result in DNA segregation^15^. For other family members, the cellular positioning of a ParA/MinD protein relies on ‘landmark’ proteins^14^. These landmark proteins serve as molecular beacons to recruit the ParA/MinD protein to specific locations in the cell. The resulting patterns are dynamic with the ParA/MinD ATPase maintaining the localization, or the other activities of the ParA/MinD protein at that site^19–22^.

The model organism *Escherichia coli*, and many rod-shaped bacteria, encode the classical example of an oscillating MinD protein. MinD in *E. coli* oscillates along the longitudinal axis of the cell, which contributes to correct placement of the Z-ring at mid-cell during cell division^23^. Consequently, deletion of the Min system leads to cell division defects, such as the occurrence of mini-cells resulting from asymmetric division^24^. In *E. coli*, MinD binds the FtsZ antagonist, MinC, and oscillates between the cell poles in a manner that results in a net higher MinCD concentration near the poles, thus preventing Z-ring assembly away from mid-cell^25^. The ATP-bound dimeric form of MinD binds the membrane via a membrane targeting sequence (MTS), while the ADP-bound monomeric form diffuses in the cytoplasm^26–28^. MinE binding to MinD induces MinD release from the cell membrane, because it stimulates the ATPase activity of MinD^10^.

ParA systems, such as the plasmid separating ParA system of *E. coli*, form dynamic patterns on the nucleoid. In contrast to the Min system, these are not simple oscillations, but instead the ParA proteins form cytoskeletal-like structures that are dynamically located over the nucleoid^29^. The best-studied ParA systems segregate plasmids and bacterial chromosomes^29–32^. Plasmid segregating ParA proteins can reversibly dimerize and bind non-specifically to the nucleoid in a nucleotide-dependent manner^13^. ParA’s ATPase stimulating factor, ParB, specifically nucleates its own binding to plasmids at the parS domain. The combined ParAB dynamics allow them to distribute plasmids over the chromosome and can effectively pull plasmids apart towards opposite cell poles. The ParA system was also found to position cytoplasmic chemotaxis proteins in *Rhodobacter sphaeroides^9^*.

Several ParA/MinD proteins have been discovered that localize without the presence of an ATPase stimulating protein, and instead bind landmark proteins to direct them to specific cellular locations^19^,^33–36^. For example, ParA/MinD homologs are responsible for the correct positioning of chemotaxis and motility machinery in several bacteria^6,37–42^.

ParA/MinD homologs are encoded in archaeal genomes^3^, but very little is known about the positioning or spatiotemporal dynamics of proteins in archaeal cells. To assess if ParA/MinD homologs in archaea play a role in cellular positioning, we selected the model archaeon *Haloferax volcanii*, which has genetic tools and fluorescent fusion proteins available to address questions relating to intracellular organization. Recent work in *H. volcanii* has indicated that positioning of the motility structure, chemosensory arrays and cell division machinery is regulated by unknown mechanisms^43–45^. *H. volcanii* cells display different shapes dependent on the growth stage of the culture^43^,^44^,^46^. Cells in early log-phase are rod-shaped and motile. They possess chemosensory arrays at the cell membrane, and filamentous motility structures, archaella, at their surface^44^. In late stationary phase, cells have a more discoid, pleomorphic appearance. They are no longer motile and do not have chemosensory arrays^44^. In this study, we identified a function of a previously uncharacterized MinD protein, designated MinD4, in *H. volcanii*. MinD4 was seen to oscillate along the long axis of the cell and is required for the correct assembly and positioning of the chemosensory arrays and the archaellum for cell motility.

## Results

### Identification of MinD-like homologs in archaea

To assess the distribution and abundance of ParA/MinD proteins in currently known archaea, we searched for homologs encoded in a diverse set of genomes representing species from across the archaeal domain. Proteins of the ParA/MinD (SIMIBI) superfamily contain the characteristic deviant Walker A motif^51^ and also include the families ArsA (arsenic/metal efflux), Mrp (membrane ion transportation), CooC (metal-enzyme incorporation), NifH (nitrogen fixation), FlhG (regulation of flagella) and others^3,14,47^. We identified at least one member of the superfamily in all archaea analyzed, and some species contained more than 12 homologs (Table S1). Phylogenetic analysis (Fig 1) showed that homologs of ParA, ArsA, Mrp and CooC were widespread across the archaeal domain. Additional groups of uncharacterized proteins from several distinct groupings as well as some distant, non-canonical sequences were also present in many species (Fig 1A, Table S1). Genes encoding MinD-like proteins were limited to the major phylum Euryarchaeota (Fig 1A, Table S1). However, the MinD-like sequences in archaea form a sister group to the bacterial MinD sequences (Fig 1B), suggesting that they diverged from a common ancestor and have evolved independently in the archaeal and bacterial lineages. We identified no specific archaeal homologs of the bacterial FlhG proteins, which are involved in positioning of the bacterial flagellum ^42^.

The *H. volcanii* genome encodes 13 genes homologous to the broader ParA/MinD (SIMIBI) superfamily (Fig 1), including four homologs of ParA, two Mrp homologs, two ArsA homologs, one non-canonical homolog, and four MinD homologs. Of the 13 homologous genes, 11 show a specific match to the deviant Walker A motif, (K/R)GGXXG(K/R). The proposed naming of the *H. volcanii* DS2 proteins in relation to their gene-identifiers is given in Fig 1B; we refer to the *minD* homologous genes as *minD1* (HVO_0225), *minD2* (HVO_0595), *minD3* (HVO_1634) and *minD4* (HVO_0322). Notably, the protein encoded by *minD4* (HVO_0322) has an unusually long C-terminal extension and shows strong homology at the very C-terminus to the membranetargeting sequence (MTS) of *E. coli* MinD (described further below). The other three *H. volcanii* MinD proteins do not have a clear MTS, even though a homologous MTS has been observed in some other more diverse MinD-like proteins from other archaea^28^. Specific homologs of MinD4, including its MTS, were identified in several other species of the *Haloferacales* order, and are well conserved in the *Haloferax* genus (Fig S6A, B).

**Figure 1:**
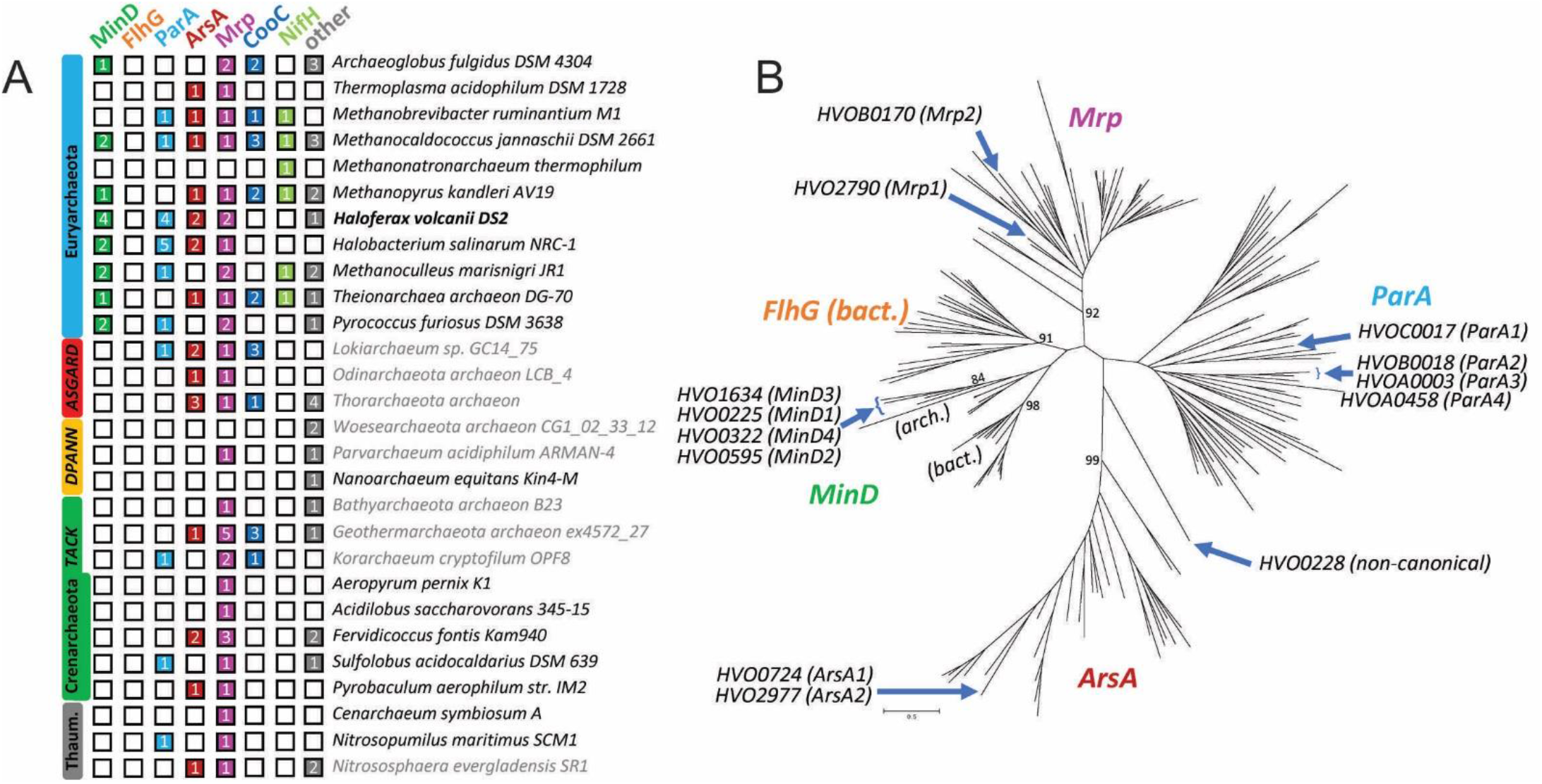
ParA/MinD-like proteins in archaea. (A) The number of protein homologs identified in the indicated archaea were determined from phylogenetic analysis. Phylum-level classification is shown on the left. The homologs include those that showed a specific match to the deviant Walker A motif. Two additional *H. volcanii* sequences that show a weaker match to the deviant Walker A motif sequence are also included (HVO0228 and HVOA0458). Species names in grey text indicate a *Candidatus* status of the taxon; in some cases, genome data may be incomplete. Sequences that cluster in groups of unknown function or those that classify weakly, or are non-canonical, are indicated in the “other” category. (B) Maximum-likelihood phylogenetic tree including curated family sequences from the MinD, ParA, FlhG, Mrp, and ArsA families and the archaeal homologs, including all 13 identified *H. volcanii* homologs. Archaea- and bacteria-specific branches are indicated. Bootstrap support for selected branches is also indicated (%).

### H. volcanii MinD homologs have no detected effect on cell shape and division

In order to identify the biological function of the four MinD homologs in *H. volcanii*, the genes encoding these proteins were individually deleted. In addition, double and triple deletion strains of several gene combinations were created, as well as a strain in which all four *minD* homologs were knocked-out. Measurement of the optical density (OD) during growth in liquid medium showed that all strains had growth rates in that were indistinguishable from the wild type (Fig 2A and S1). Next, all strains were observed by light microscopy and the cell size and shape was analyzed by determining the cell circularity (cell elongation) and area. Cell volume distributions were also obtained by Coulter cytometry, as a sensitive assay for defects in cell division. These analyses revealed no cell size or shape differences compared to wild-type in any of the *minD* mutant strains (Fig 2B-E and Fig S2). Furthermore, to assess if the four *H. volcanii* MinD homologs had any effect on positioning of the Z-ring, fluorescence microscopy was used to visualize localization of FtsZ1-GFP in *minD* knock out strains (Fig 2F, G). In all single deletion strains, as well as the quadruple-deletion strain, the FtsZ ring was still positioned correctly at mid-cell, similar to the FtsZ1-GFP localization pattern in wild type *H. volcanii* (Fig 2F, G, S3). These results suggested that, under the conditions investigated, the four MinD proteins are not important for cell division or cell shape in *H. volcanii*.

**Figure 2:**
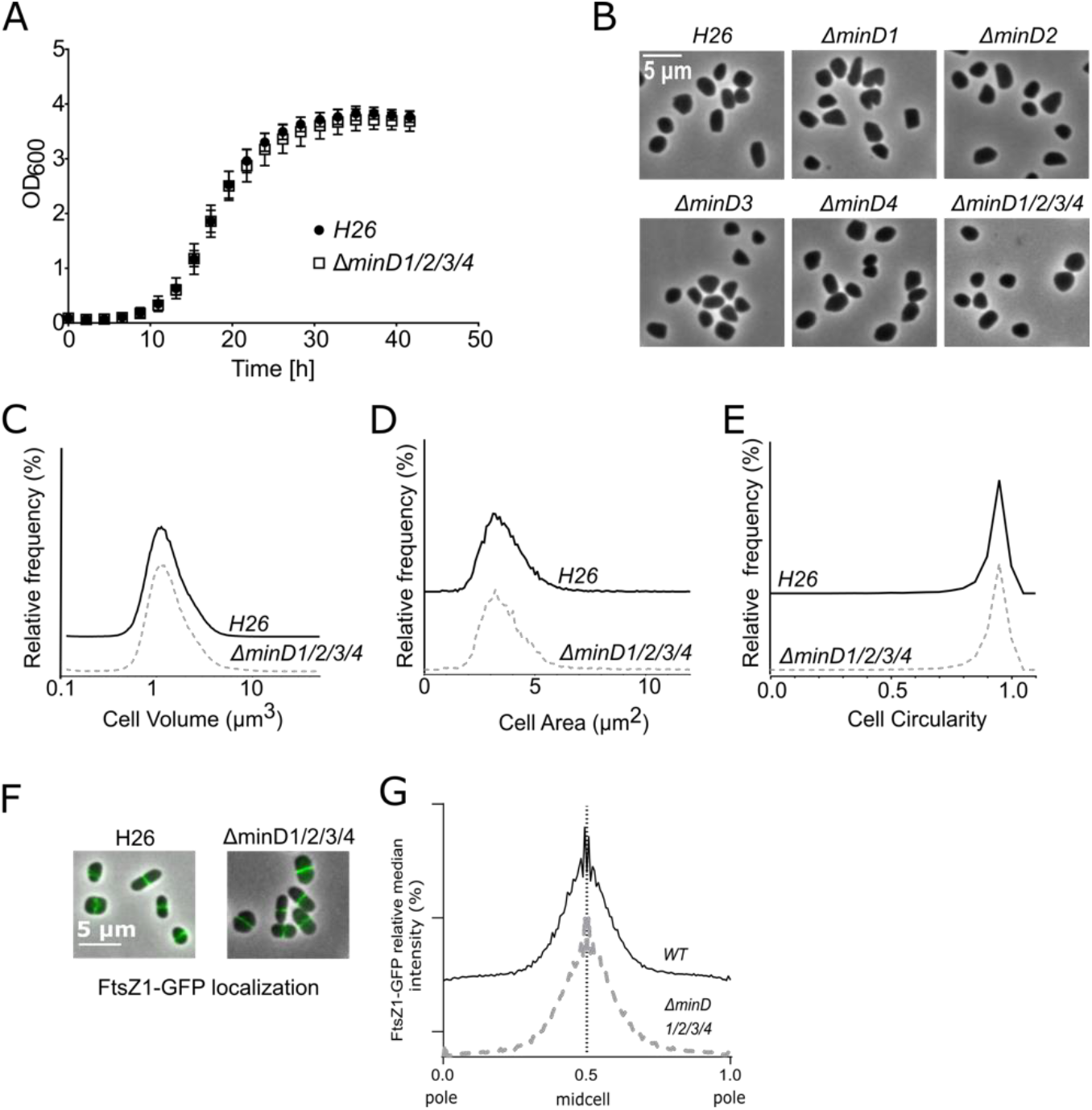
Archaeal mind genes do not affect cell shape and Z-ring I positioning. (A) Optical Density measurements in CAB medium at °45 C of a *H. volcanii* strain deleted for all four minD genes, compared to wild type *H. volcanii* (H26) cells. (B to G) *H. volcanii* shape analysis in exponential phase in CAb medium. (B) Phase contrast images of *H. volcanii*. (C) Relative frequency distributions of cell volume measured by Coulter cytometry n>200000. (D) Relative frequency distributions by automated analysis of cell area from cells analyzed in B. (E) Relative frequency distributions by automated analysis of cell circularity from cells analyzed in (B). All results are the average of at least two representative independent experiments. Sample sizes for (D) and (E): n_H26(WT)_ =5730, n*_ΔminDi/2/3/4_*=10930. (F) FtsZ1-GFF localization in quadruple *minD* mutants during exponential growth phase. Figures show an overlay of phase contrast and GFP fluorescence channels. (G) Representation of the relative median intensity of the FtsZ1-GFP signal along the relative cell length measured by automated image analysis. Sample sizes: n_WT(H26)_ = 1252, n*_ΔminD1/2/3/4_*=2747.

### MinD4 is important for motility

The *minD4* gene is located near genes encoding the motility and chemotaxis machinery (*flaD* and *cheW*) in several related Haloarchaea (such as *Haloarcula* sp.). In addition, the STRING database indicated that *minD4* is in the genomic neighborhood of the chemotaxis genes *cheY, cheR* and *cheB* in several haloarchaeal genomes^48^. Thus, we hypothesized that MinD4 might be involved in motility and chemotaxis instead of cell division. We tested the motility behavior of the Δ*minD4* strain on a semi-solid agar plate, which allows cells to swim out from the original inoculation site and form a ‘motility ring’. Directional movement requires functional motility machinery and a signal-sensing taxis system. After 4-5 days of incubation at 45 °C, wild-type *H. volcanii* forms motility rings of several centimeters across. The Δ*minD4* strain was still able to form a motility ring, however, the diameter was reduced by ~50% in comparison to the wild-type strain (Fig 3). Hence, MinD4 is important for directional movement and might affect motility and/or chemotaxis machinery.

**Figure 3:**
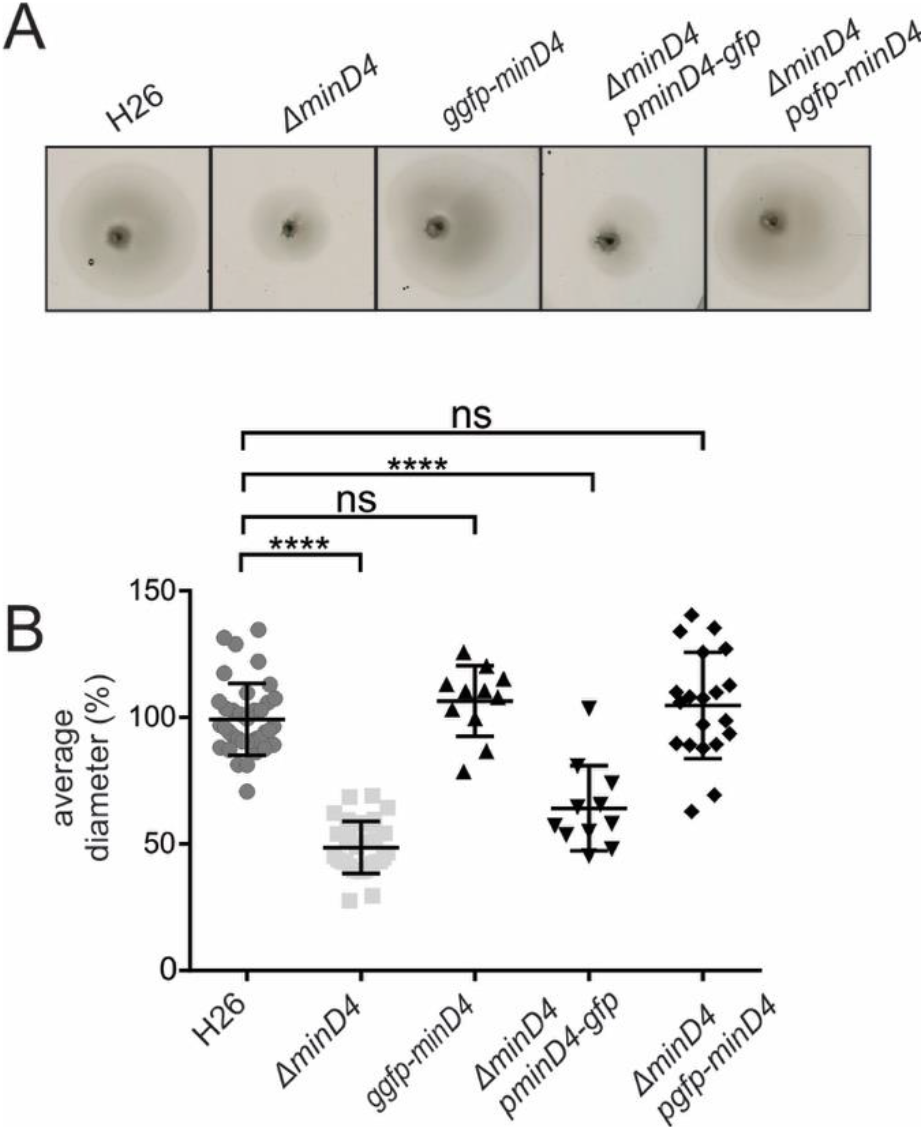
Effect of *minD4* deletion on motility. (A) Representative examples of motility assays on semi-solid agar plates of different *H. volcanii* strains, indicating that deletion of *minD4* leads to reduced directional movement. (B) Average diameter of motility rings, measured relative to the wild type, from different *H. volcanii* strains from >3 independent experiments including 4 biological replicates each. Middle black line indicates mean, lower and upper lines the standard deviation. gGFP, genomic integration of gfp-minD4.

### MinD4 displays polar localization and oscillation behaviour in cells

To gain more insight into the biological function of MinD4, we decided to study its cellular positioning. Both N-terminal and C-terminal GFP fusions to MinD4 were expressed under the tryptophan inducible promotor in a *ΔminD4* strain. Correct expression was confirmed with western blotting, which showed a band around 150 kDa, the size of a dimer (Fig S4). Expression of the N-terminal (GFP-MinD4) fusion restored the motility phenotype of the Δ*minD4* strain on tryptophan containing semi-solid agar plates, demonstrating that the fusion protein was functionally indistinguishable from the wild-type protein in this assay, and that the Δ*minD4* motility phenotype can be successfully complemented by reintroduction of a *minD4* ORF (Fig 3). We also generated a strain with a genomic copy of GFP-MinD4 inserted at the *minD4* locus with the native *minD4* promoter and found this was also fully functional (Fig 3). On the other hand, the C-terminal (MinD4-GFP) fusion could not restore motility (Fig 3), and we therefore continued with fluorescence microscopy studies using the GFP-MinD4 fusion as the only source of MinD4 in the cells.

We first observed that the GFP-MinD4 signal in cells in the exponential growth phase was most intense at one or both cell poles, as a focus or distinct patch-like structure. The positioning pattern of the GFP-MinD4 protein was similar when expressed from the genome (gGFP-MinD4), or from a plasmid (pGFP-MinD4) under the control of the tryptophan-inducible promoter, *p.tna* (Fig 4A, B). As deletion of minD4 had an effect on motility, we also aimed to study its cellular positioning in motile cells. Observation of rod-shaped motile cells sampled from the early-log phase (Fig 4C) showed a similar positioning to that of non-motile cells from exponential phase (Fig 4B). The GFP-MinD4 signal was also most intense at both cell poles, where it formed patch-like structures. In addition, we observed that in relatively long cells, GFP-MinD4 also appeared at mid-cell (Fig 4C, D), suggesting that part of the MinD4 proteins concentrates at mid-cell as cells become larger or prepare to divide.

**Figure 4:**
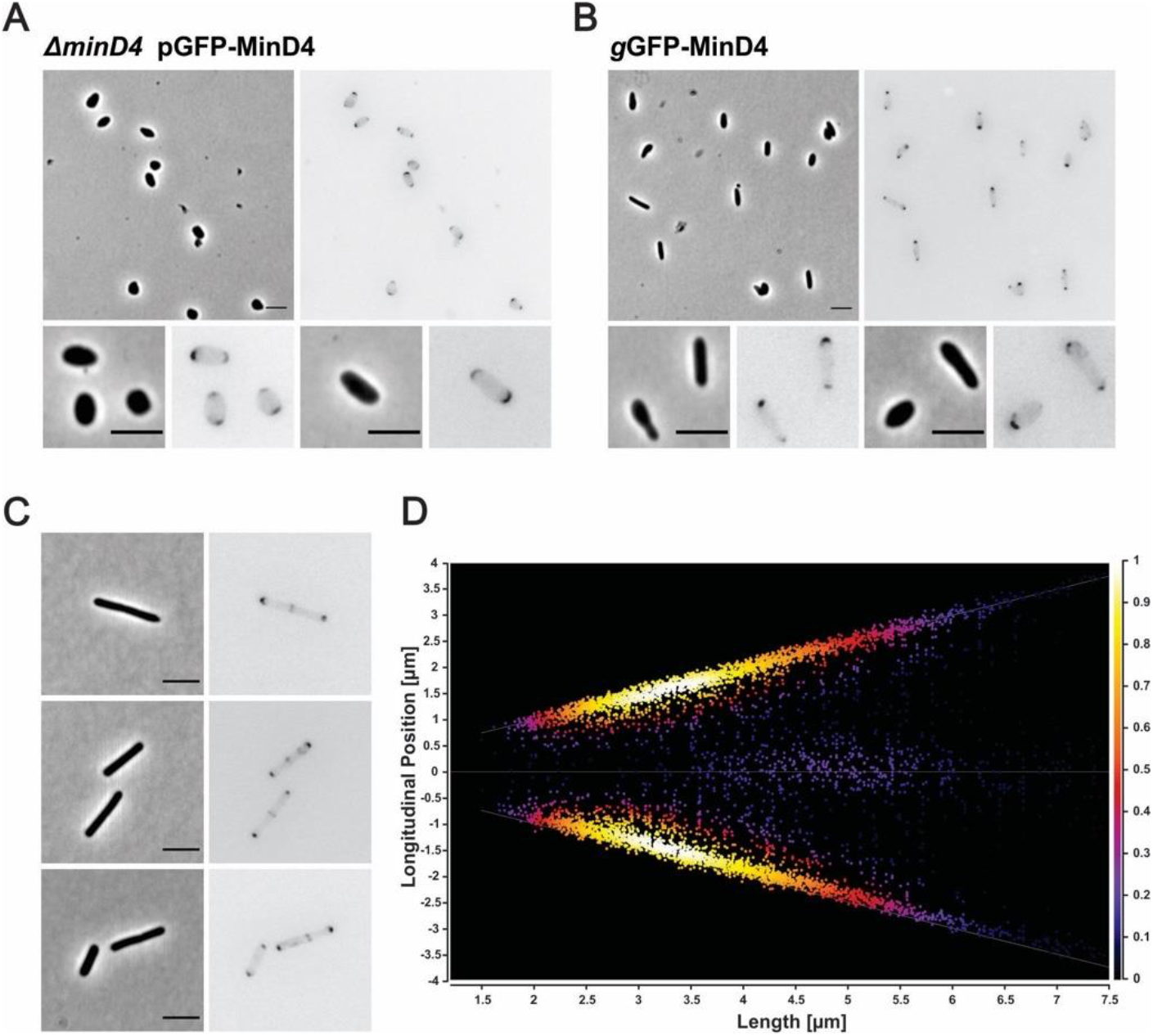
Cellular positioning of MinD4. (A) Fluorescent images of GFP-minD4 localization in the Δ*minD4* strain or (B) by genomic integration of an N-terminal fusion of GFP to minD4 (gGFP minD4) of *H. volcanii* cells in exponential phase. Scale bar 4 μm. (C) Representative fluorescence images of cells expressing gGFP-MinD4 at early log phase (OD 0.05). Scale bar 4 μm (D) Cellular distribution of MinD4 signal. The signal along the longitudinal cell axis is shown at the Y-axis, where 0 indicates mid-cell. Cells are arranged according to their size, with shorter cells appearing on the left and longer cells on the right of the figure. Colours indicate the intensity of the signal, where white is the most intense signal. In addition to polar signals, longer cells also show signal at mid-cell (N>2000).

The distribution of MinD4 comprises bright foci located at the cell poles as well as a much dimmer continuous distribution of MinD4 throughout the cell (Fig 4A, B). Interestingly, time-lapse imaging revealed that the background distribution of MinD4 oscillates along the long axis of cells (Fig 5A-E). The large foci observed at cell poles remain effectively stationary inside cells during oscillations, although their intensity changes during the period of oscillation (Fig 5E). The strain carrying the genomic GFP-MinD4 showed oscillation with a regular period of approximately 210 seconds (Fig 5A, D, Movie S1). Similarly, GFP-MinD4 expressed from a plasmid had a slightly faster oscillation with an average period of 160 seconds (Fig 5A, E) with a similar relative amplitude distribution (the amplitude of oscillation divided by the total fluorescence in the cell) (Fig 5B). Two different patterning regimes were observed. Motile, rod-shaped cells from exponential phase generally oscillated pole to pole similar to the *E. coli* Min protein system (Fig 5C, D, E, Movie S2). Round/plate-shaped cells sometimes displayed a different type of patterning where the maxima rotated around the edge of the cell (Fig 5F, Movie S3). This is similar to patterning observed in the *E. coli* Min system when it is confined in artificial square shapes^49^. Thus, a background distribution of MinD4 displays oscillatory movement inside cells, which has some similarity with the oscillatory behavior of MinD from *E. coli*. However, MinD4 in addition forms static polar patches that are not observed in the *E. coli* system. We then investigated whether deletion of the *minD1*, *minD2* or *minD3* genes would affect the positioning pattern and oscillation behaviour of GFP-MinD4. Similar to in the Δ*minD4* background, MinD4-GFP was positioned at the curved membranes near the cell poles in the Δ*minD1*, Δ*minD2* and Δ*minD3* strains (Fig S5A). In addition, oscillation of the background distribution of MinD4 still occurred in the three deletion backgrounds as is the case in the Δ*minD4* strain (Fig S5B). In conclusion, MinD1, MinD2 and MinD3 are not individually required for subcellular localization of MinD4.

**Figure 5:**
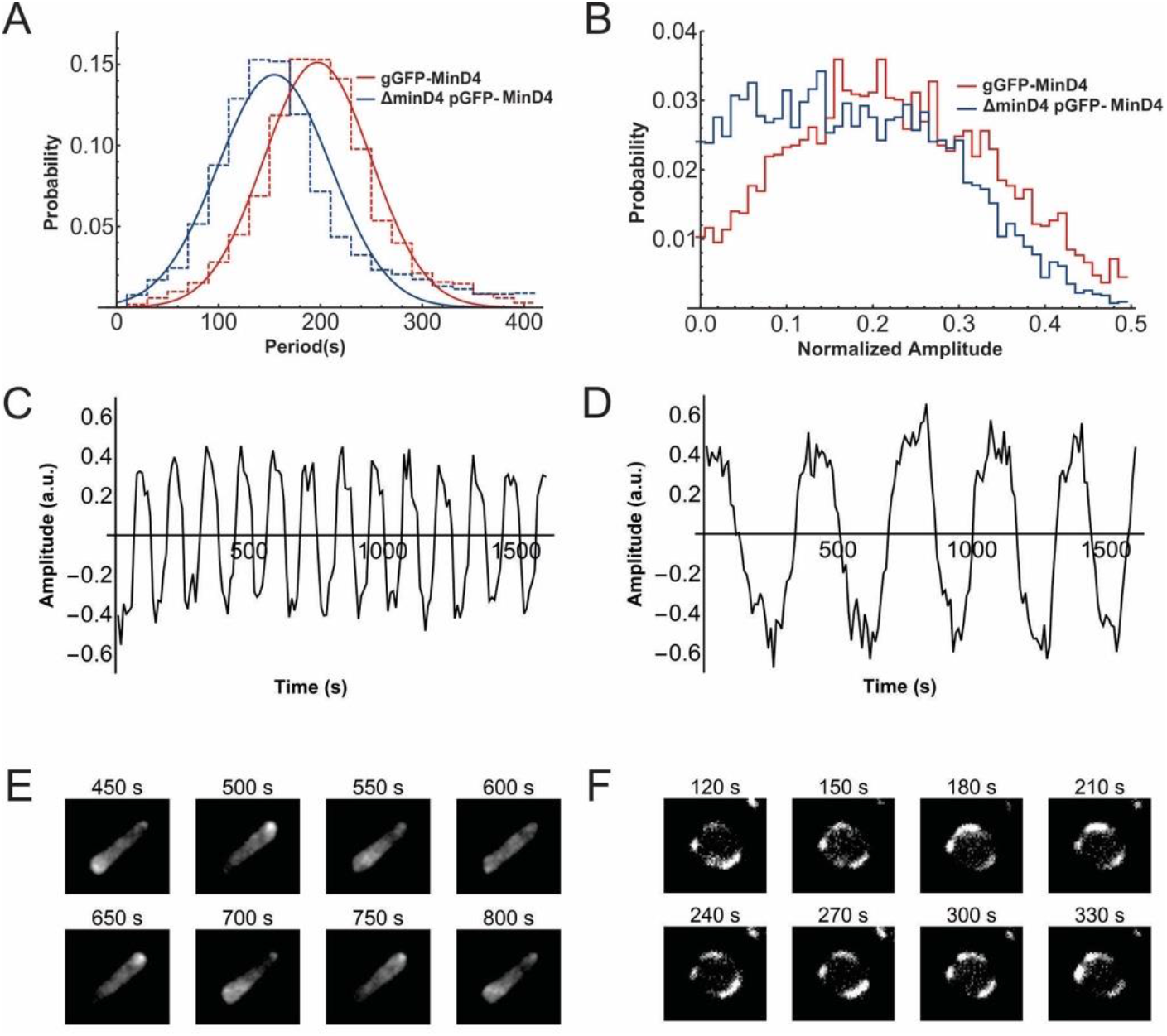
Oscillations of MinD4 Patterning. (A) The distribution of periods of oscillations observed for each variant of MinD4. (B) The relative amplitude distribution for each variant of MinD4 given by the amplitude of oscillations divided by the mean fluorescence within the cell. (C) A trace showing the typical oscillation of the ΔminD4 pGFP-MinD4 strain expressing GFP-MinD4 from plasmid in rod like cells. (D) A trace showing the typical oscillation of the genomically expressed GFP-MinD4 (gGFP-MinD4) in rod like cells. (E) Images of corresponding Δ*minD4* rod-shaped cell showing GFP-MinD4 oscillation from pole to pole in the cell. (F) Images of GFP-MinD4 in a Δ*minD4* round cell which goes around in a circle rather than from pole to pole.

### Importance of ATP Hydrolysis for MinD4 functionality

MinD4 contains a Walker A and a Walker B motif that are critical for ATP binding and hydrolysis in bacterial MinD homologs. To investigate the role of ATP hydrolysis in the function of MinD4, we mutated the deviant Walker A motif (K16A) and Walker B motif (D117A) and observed the motility behavior of the mutants (GFP-minD4_WA* and GFP-minD4_WB*) expressed from a plasmid in the Δ*minD4* strain on semi-solid agar plate. The motility rings were reduced to ~50% of wild type, just as for the full *minD4* deletion strain (Fig 6A). Both mutated fusion proteins were correctly expressed as judged by Western blot (Fig S4). The subcellular localization of GFP-minD4_WA* and GFP-minD4_WB* differed substantially compared to the wild-type GFP-MinD4. The most significant difference was the lack of the bright polar patches and foci in the mutants (Fig 6B). However, gradients of fluorescence intensity were still observed along the long axis in many cells (Fig 6B), and, surprisingly, the GFP-minD4_WA* and GFP-minD4_WB* mutants still oscillated along the cell axis (Fig 6C-H, Movie S4, S5). GFPminD4_WA* oscillated with close to the same mean period as GFP-MinD4 (140 seconds versus 160 seconds respectively (Fig 6C, E)). The overall relative amplitude (fluorescence intensity) of oscillation of the GFP-minD4_WA* mutant was decreased relative to the wild type proteins due the lack of bright foci (Fig 6D, E, G). Similar as for the WalkerA mutant, the GFP-minD4_WB* oscillation period was comparable to wild type MinD4 and there was also a decreased amplitude (Fig 6C, D, F, H Movie S5). As ATP hydrolysis is critical for the function of ParA/MinD like proteins, it appears that MinD4 ATP hydrolysis is not powering the oscillation pattern but rather is required for the formation of the polar patches. This phenotype is similar to Par proteins^15^.

**Figure 6:**
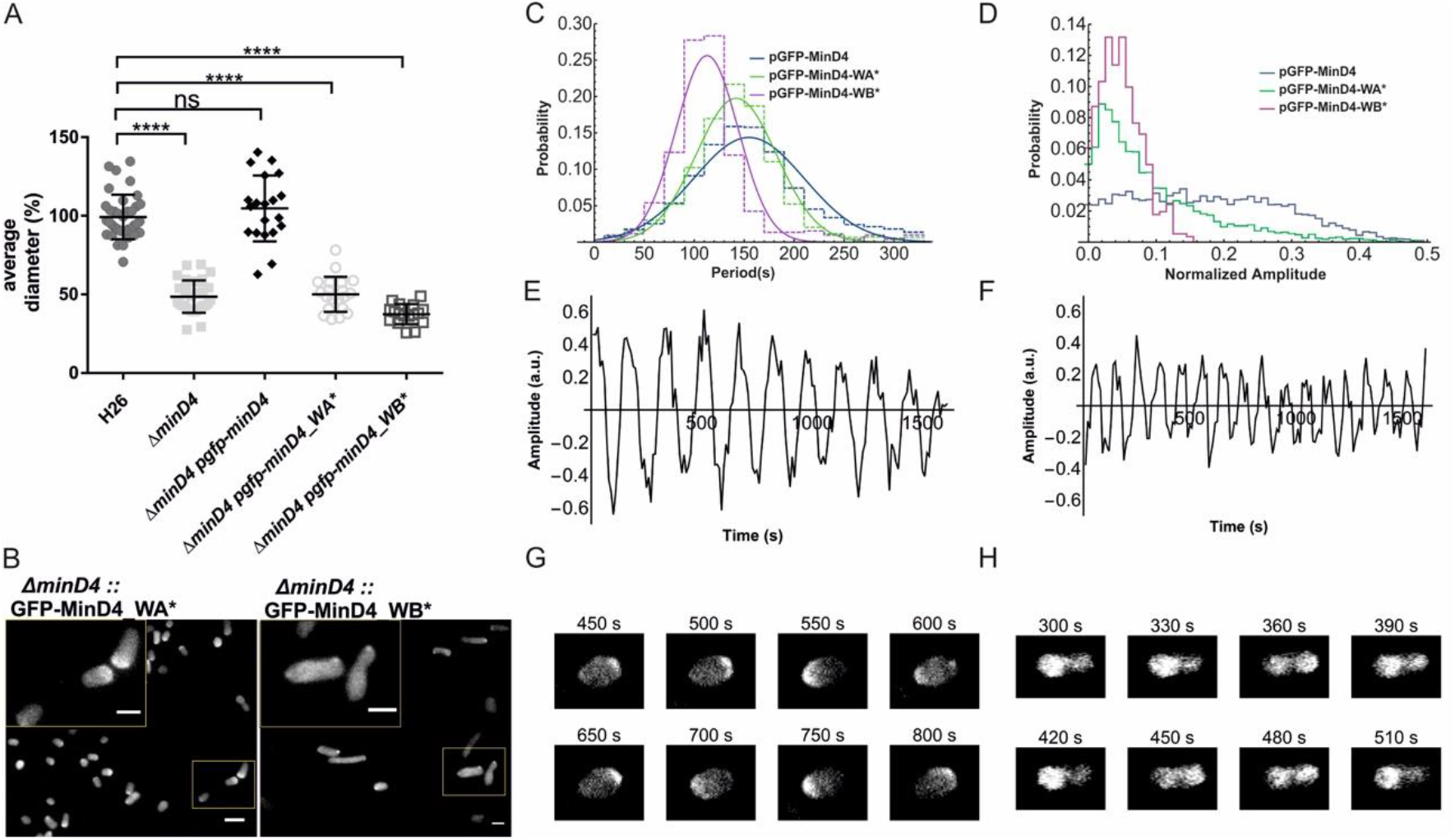
Formation of polar patches by MinD4 relies on the deviant WalkerA and WalkerB motif. (A) Motility on semisolid agar plate is affected by mutation of the WalkerA (K16A, MinD4_WA*) and WalkerB (D117A, MinD4_WB*) motif. Average diameter of motility rings, measured relative to the wild type, from different *H. volcanii* strains from >3 independent experiments including 4 biological replicates each. Middle black line indicates mean, lower and upper lines the standard deviation. (B) Fluorescent images of a ΔFluorescent images of a*minD4* pGFP-MinD4_WA* (left) and pGFP-MinD4_WB* (right). Scale bar large panel, 4 μm. Scale bar inset, 2 μm. (C) The distribution of periods of oscillations observed for each variant of MinD4. (D) The relative amplitude distribution for each variant of MinD4 given by the amplitude of oscillations divided by the total fluorescence within the cell. (E) A trace showing the typical oscillation of the Δ*minD4 pGFP-minD4_WA\** in rod shaped cells. (F) A trace showing the typical oscillation of the Δ*minD4 pGFP-minD4_WB\** in rod shaped cells. (G) Images of a Δ*minD4 cell* expressing *pGFP-minD4_WA\** cell showing that this mutant still oscillates but does not display the polar patches seen in the wild type. (H) Images of a Δ*minD4* cell expressing *pGFP-minD4_WB\** showing that this mutant still oscillates but does not display the polar patches seen in the wild type.

These results also suggest that MinD4 is likely binding to an alternative patterning system to give rise to the observed oscillations of the protein.

### The membrane targeting sequence is essential for MinD4 function

Members of the bacterial MinD family can have a membrane targeting sequence (MTS) at their C-termini. This MTS forms an amphipathic helix allowing membrane interaction and was found to be important for localization of MinD in *E. coli* and *B. subtilis*, and FlhG in *Geobacillus thermodenitrificans*^26,27,42,50^. Protein sequence alignments of the C-termini of bacterial MinDs and FlhGs revealed that MinD4 has a *bona fide* MTS at their extreme C-termini, consisting of ~7 amino acids, which are predicted to form an alpha helix (Fig 7A, S6A, B)^28^,^51^. Interestingly, an alignment of MinD4 homologues showed that the conservation of the C-termini is not restricted to the MTS but spans along a stretch of ~40 amino acids, referred in this study to as extended MTS (eMTS). This region is predicted to contain a second alpha-helix (Fig. 7A, S6B). In the long C-terminal extension of MinD4, between the ATPase domain and eMTS, several blocks rich in aliphatic and acidic amino acids, are present (Fig S6B).

**Figure 7:**
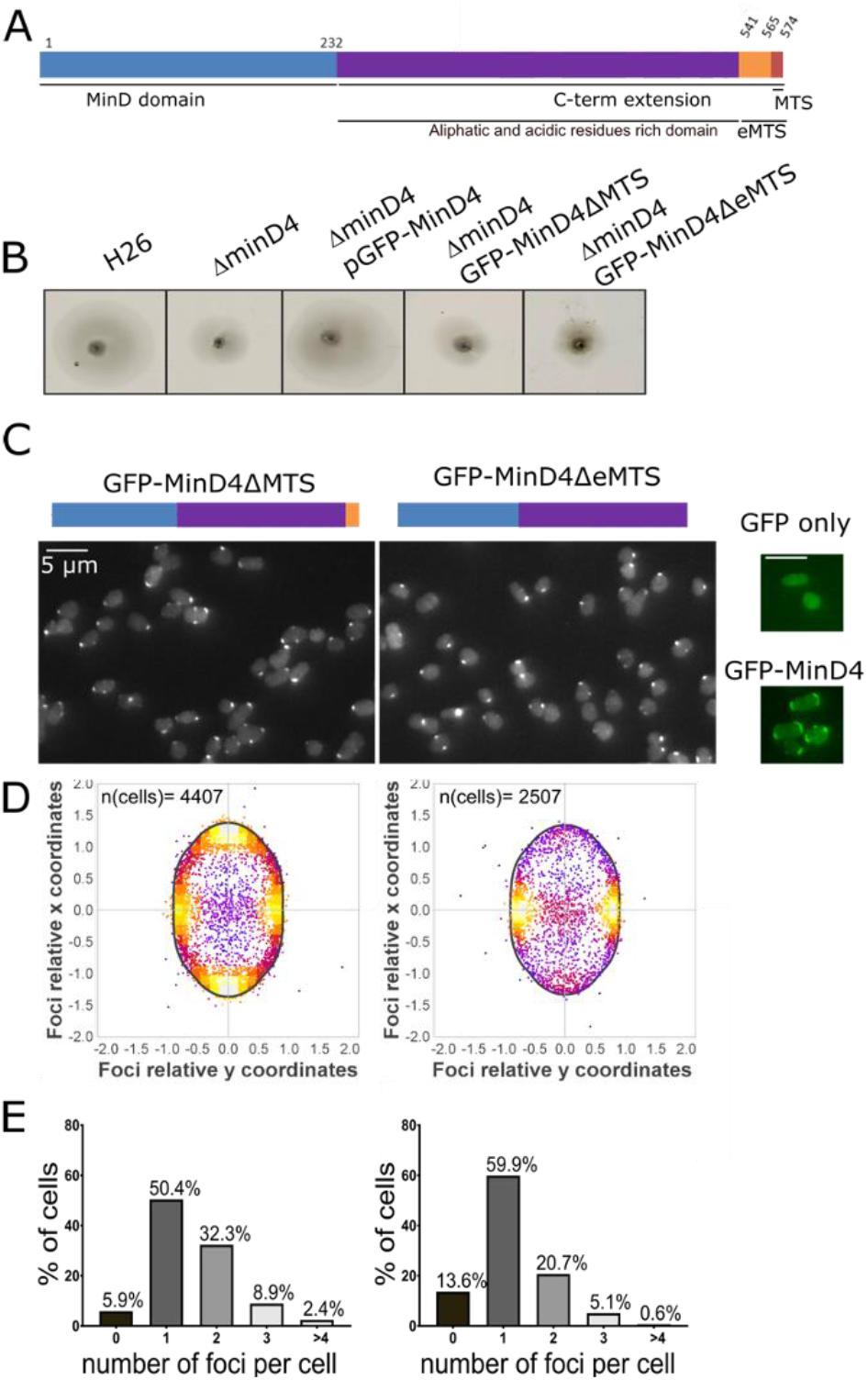
C-terminal domains of MinD4 are necessary for its correct localization in *H. volcanii*. (A) Schematic representation of MinD4 protein domains. The numbers indicate the number of the amino acids based on *H. volcanii* MinD4 sequence. (B) Effect of C terminal protein truncations on MinD4 localization in *H. volcanii*. (C) Schematic representation of the truncated GFP-MinD4 proteins and the corresponding cellular localization studied in Δ*minD4* by fluorescence microscopy. GFP fluorescence channels are showed either in green or grey. GFP only: Δ*minD4* + empty GFP expressing vector, GFP-MinD4ΔMTS and GFP-MinD4ΔeMTS: 0.6 sec at 50% intensity. (D) Density map of the relative position of GFP-MinD4 truncations foci in relation to the cell shape analysed by automated image analysis. The vertical line indicates the longitudinal cell-axis and bold black line represent an idealized oblong cell shaped. (E) Histogram of the percentage of cells as a function of the number of GFP-MinD4 foci detected per cell analysed by automated image analysis. All samples were grown in CAB medium and sampled in logarithmic growth phase.

To assess the role of the C-terminal region for the function of MinD4, truncations were created: MinD4ΔMTS (1-560), MinD4ΔeMTS (1-535) and MinD4ΔCterm (1-232). These truncated proteins were fused to GFP at their N-termini and their expected expression was confirmed by Western blot (Fig S4). Only the MinD4ΔCterm showed protein signal at an unexpected position. This indicated that this truncated protein was not correctly expressed (Fig S4). We therefore focused mainly on the two other MTS mutants. We tested whether these mutants could complement the motility defect observed in a *ΔminD4* background. All truncation mutants were non-functional in motility assays, as they showed motility-ring diameters that were indistinguishable from the *ΔminD4* strain (Fig 7B S6C). Thus, the MTS is important for the biological functioning of MinD4.

Next, the cellular localization patterns of the truncated MinD4 proteins were studied in exponentially grown cells. The MinD4ΔMTS and MinD4ΔeMTS mutants generally localized as foci around the cell edges, which differed substantially from the typical positioning of the wild-type protein as patches at the poles (Fig 7C). Strains expressing the MTS or eMTS truncations showed one or more foci in the majority of cells (94% and 86% for ΔMTS and ΔeMTS, respectively) (Fig 7E).

We then analyzed the foci radial positioning relative to the long axis of cells (Fig 7D). The foci observed in cells expressing MinD4ΔMTS were positioned at the cell poles or at mid cell, and those of the MinD4ΔeMTS GFP fusions, were found exclusively at mid-cell (Fig 7D). Importantly, all mutants of the MTS of MinD4 displayed effectively no oscillating background distribution, as seen for wild type MinD4 (Fig S6D, E, Movie S6, S7). The above findings suggest that the eMTS is important for the oscillation and the formation of polar patches of MinD4.

### MinD4 influences cellular positioning of motility and chemotaxis machinery

The above analysis of the cellular localization of MinD4 in archaea by fluorescent microscopy, indicates that the protein behaves in a manner somewhat similar to *E. coli* MinD, as it oscillates along the cell’s long axis However, since cell division and cell shape of *H. volcanii* were not affected by *minD4* deletion, the function of *H. volcanii* MinD4 is not identical to *E. coli* MinD. Because the *H. volcanii* Δ*minD4* strain has a motility defect, we hypothesized that MinD4 might have a similar function to other ParA/MinD homologs from several bacterial species, which are involved in the placement of the flagellum and chemotaxis machinery^6,9,14,52^.

First, to determine whether the *minD4* deletion influenced formation of the archaeal motility structure, the archaellum, transmission electron microscopy (TEM) of negatively stained cells was performed. A *ΔpilB3* background strain, lacking pili, facilitated the correct identification of archaella. Electron micrographs of Δ*pilB3*Δ*minD4* showed that these cells displayed archaella at the surface (Fig 8A). But the percentage of cells with archaella was significantly reduced in comparison with the wild-type strain (Fig 8B).

**Figure 8:**
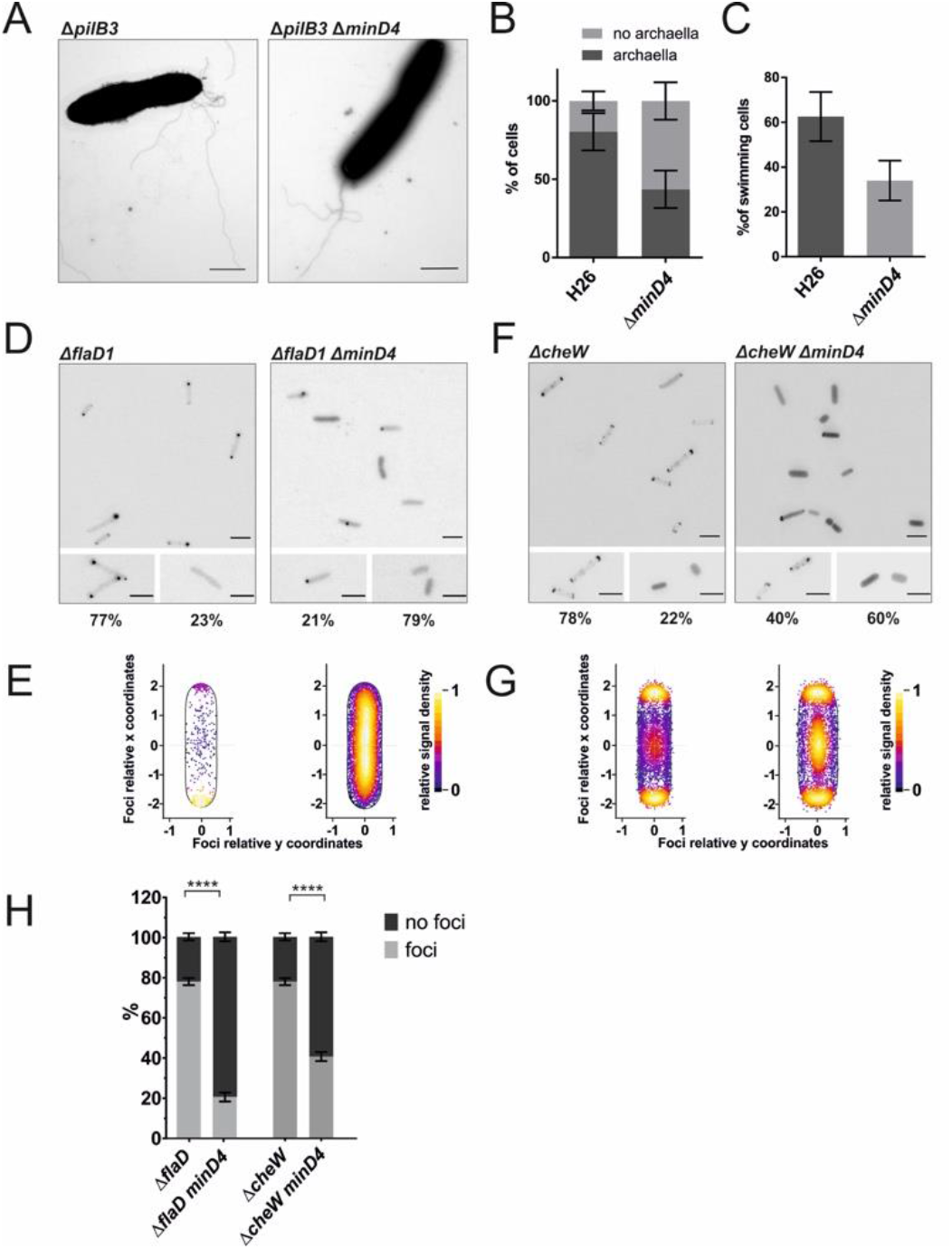
Positioning of chemosensory arrays is affected in *H. volcanii ΔminD4*. (A) Transmission electron microscopy of Δ*pilB3* and Δ*pilB3*Δ*minD4 H. volcanii* cells in early exponential growth phase. Scale bar, 1 μm. gGFP, genomic integration of gfp-minD4. (B) Percentage of cells with or without archaella filaments at the surface, as analysed by TEM. Control, n=51, Δ*minD4*, n=61 (C) Percentage of swimming cells as analysed by time-lapse imaging of wild type and Δ*minD4 H. volcanii* strains. Experiment was performed on 5 independent occasions. For each strain, 3-10 time lapse images containing ~50 cells each were analysed. The number of cells displaying clear swimming motility was divided by the total number of cells visible in the first frame of the time lapse movie. (D) Exemplary fluorescent images of intracellular distribution of FlaD-GFP in *ΔflaD* or *ΔflaD ΔminD4* expressing pFlaD-GFP from plasmid in *H. volcanii* in the early log phase (OD_600_ 0.01). The lower frames are a close-up of two different distribution patterns. The numbers below represent the percentage of the total population displaying this distribution. Scale bars 4 μm (E) Distribution of intracellular FlaD or foci. Left: pFlaD-GFP in *ΔflaD*, Right: pFlaD-GFP in *ΔflaD ΔminD4*. The cluster distances were plotted as a percentage of the total cell length. (N>2000). (F) Exemplary fluorescent images of intracellular distribution of pGFP-CheW expression from plasmid in *ΔcheW* or *ΔcheW ΔminD4*. Scale bars 4 μm. (G) Distribution of intracellular CheW foci. Left: pGFP-CheW in *ΔcheW*, Right: pGFP-CheW in *ΔcheW* Δ*minD4*. The cluster distances were plotted as a percentage of the total cell length. (N>2000). (H) Distribution of CheW or FlaD localization (green) and diffuse signal (blue) in *H. volcanii* at different growth stages.**** p< 0.0001, * p<0.05.

To assess the swimming behavior of this mutant, we imaged live cells over time in liquid medium. Cells were serially diluted and observed at early log-phase. *H. volcanii* wild type cells are motile and rod-shaped in early log phase and round and non-motile in stationary phase^44^,^46^. The majority of observed cells of the wild-type performed a random walk in the absence of directional stimuli, as has been observed before^46^ (Movie S8). Cells of the Δ*minD4* strain also performed a random walk. However, a significantly lower number of swimming cells were observed in comparison with the wild-type (Fig 8C, Movie S9). Thus, deletion of *minD4* leads to a reduced number of motile and archaellated cells.

To allow for an independent quantitative analysis of the number of cells with archaella, we used fluorescence microscopy. We selected FlaD as a well-established marker protein for the archaellum^6,44^. The FlaD-GFP fusion was previously shown to be fully functional^44^. The positioning of FlaD in the archaellum, is schematically shown in Fig S7A.

We observed *H. volcanii* cells in early log phase expressing FlaD-GFP in a Δ*flaD* or a Δ*flaDΔminD4* background. In wild type cells, FlaD foci were found exclusively at the cell poles in the majority of cells (~80%)(Fig8D, E), as it was previously shown that *H. volcanii* has polar bundles of archaella ^44^. When minD4 was deleted, the number of cells with FlaD foci, was reduced significantly to ~20% (Fig 8D, E, H). The remaining cells showed diffuse fluorescence in the cytoplasm, indicating that no archaellum motors are formed in these cells. These findings are in correspondence with the TEM analysis of a Δ*minD4* strain that showed a reduced number of cells with archaella (Fig 8B). FlaD-GFP expression was also studied at higher ODs, when cells are no longer motile. In this case FlaD foci are present in all cells, and were previously hypothesized to be remnants of the archaellum motor^44^. Consequently, the observed difference between the wild type and minD4 deleted strain is most obvious in early exponential phase (Fig S7B).

Next, we focused our attention to the positioning of the chemosensory arrays in *H. volcanii*. The CheW protein was previously demonstrated to be a good marker protein to study the positioning of chemosensory arrays in *H. volcanii*, and a GFP fusion is fully functional^44^. The localization of GFP-CheW was studied in a *ΔcheW* strain and a *ΔcheWΔminD4* strain. Normally, the chemosensory clusters of *H. volcanii* have a polar preference in rod-shaped motile cells in early log-phase^44^. In addition, smaller clusters are observed along the lateral membranes^44^. Consistent with this, GFP-CheW in cells in early-log phase of the Δ*cheW* strain showed foci near the cell poles and along the lateral membranes in ~80% of the cells (Fig 8F, G, H). In the remaining ~20%, the GFP-CheW signal was diffuse in the cytoplasm, suggesting that these cells have no detectable chemosensory arrays (Fig 8F, H). In contrast, in the Δ*cheW*Δ*minD*4 strain, distinct GFP-CheW foci at the membrane were only observed in 40% of cells (Fig 8 F,G, H). In the other 60% of cells, the GFP-CheW signal was diffuse in the cytoplasm (Fig 8H), suggesting that chemosensory arrays are not present. In those *ΔminD4* cells where distinct CheW foci were observed, the positioning pattern was similar to that observed for wildtype cells (Fig 8F).

Previously it was shown that chemosensory arrays dismantle when the cell culture reaches stationary phase and most cells are round and non-motile. To analyze whether chemosensory arrays are absent in stationary phase in the Δ*cheW*Δ*minD*4 strain, we studied GFP-MinD4 in this strain at different growth phases (Fig S7C). This revealed that deletion of minD4 resulted in a significantly lower percentage of cells with chemosensory arrays in exponentially growing cells (until OD 0.6). In stationary phase, cells become discoid-shaped, are no longer motile, and both the *ΔcheW* and a *ΔcheWΔminD4* strain expressing GFP-CheW display diffuse fluorescence, suggesting that chemosensory arrays are absent (Fig S7C). As the number of cells with chemosensory arrays is already lower at early exponential phase, the overall cell population of the ΔminD4 strain is devoid of chemosensory arrays earlier during growth when compared to the wild type (Fig S7C). Thus, in the Δ*cheW*Δ*minD*4 strain, the number of cells with chemosensory arrays is reduced, indicating that MinD4 promotes the assembly of chemosensory arrays in *H. volcanii*.

In wild-type cells, both lateral and polar chemosensory arrays are mobile within the membrane^44^. We applied time-lapse imaging of Δ*cheW*Δ*minD4* cells expressing GFP-CheW and found that the chemosensory arrays in this strain showed similar mobility patterns compared to the wild-type strain (Fig S7D, Movie S10); polar clusters mainly remained in the polar region, while lateral clusters moved freely along the lateral membranes. (Fig S7D Movie S10). Next, we followed the movement and new generation of chemosensory arrays during growth and cell division over 16 hours in cells of the Δ*cheW*Δ*minD4* strain. We occasionally observed the birth of new chemosensory arrays (Fig S7E), similar as the wild type cells. Still, eventually the total number of cells with chemosensory arrays is reduced in comparison with the wild type.

Together these results suggest that those Δ*minD4* cells that do manage to form the motility structure and chemosensory arrays, still present those structures in the correct cellular position. However, deletion of *minD4* leads to a significant reduction of both the number of cells with archaella and with chemosensory arrays.

## Discussion

ParA/MinD proteins are involved in the positioning of macromolecular protein complexes in bacteria. Genes encoding ParA/MinD homologs are also distributed widely, but not universally, amongst the archaeal domain. Although members of almost all archaeal phyla encode ParA homologs, MinD homologs are exclusively found amongst the Euryarchaeota. The crystal structures of the MinD proteins from the euryarchaea *Pyrococcus horikoshii, Pyrococcus furiosus* and *Archaeoglobus fulgidus* were solved and show a classical MinD fold, however, these proteins are monomeric in the crystalsirrespective of the bound nucleotide^53–55^. *In vitro* analysis of the *P. horikoshii* MinD revealed a low level of ATPase activity in comparison with other motor proteins^53^. Besides this initial structural analysis, no further characterization of the biological function of MinD in archaea was attempted until now. Interestingly, the archaeal MinD-like sequences form a separate group from the bacterial MinDs, which suggested that they might have a different function than bacterial MinDs. We aimed to address the biological function of MinD proteins in archaea using the model archeon *H. volcanii*. We found that the four MinD homologs encoded by *H. volcanii* are not involved in cell division, in contrast to bacterial MinDs. Instead, one of the homologs, MinD4, was shown to affect motility, due to its role in the cellular positioning of the motility and chemotaxis machinery, which are required for directional movement.

Several orphan ParA/MinD homologs were reported to be involved in positioning of flagella and chemosensory arrays in bacteria. For example, a ParA homolog, designated ParC, is required for correct placement of clusters of chemotaxis receptors at the cell poles of *Vibrio sp*. by binding to the landmark protein HubP^6^,^20^,^37^. In addition, the MinD homolog, FlhG, is responsible for maintenance of the position and number of flagella in a wide range of bacteria, such as in *Vibrio sp, Pseudomonas aeruginosa*, *Shewanella putrefaciens, Heliobactor pylori* and *Bacillus subtilis*^38–42^. As FlhG has a MTS, similar to other MinD like proteins, the dimeric state is targeted to the membrane^42^. Archaea do not possess an FlhG homolog (Fig 1). All above examples function in concert with a landmark protein, which targets them to specific cellular positions. Therefore, these proteins do not oscillate, but instead remain static in the cell. A possible exception is a parA homolog that positions the large cytoplasmic chemosensory array from *Rhodobacter sphaeroides* at mid cell, and was suggested to oscillate on the nucleoid^9^.

Fluorescence microscopy of GFP tagged MinD4, showed that, in addition to polar patch formation, the protein can oscillate along the length of rod-shaped *H. volcanii* cells (Fig 9). The formation of polar patches is dependent on ATP binding and hydrolysis as mutation of WalkerA or WalkerB motifs abolishes the formation of these patches. However, the oscillation of the background distribution of MinD4 is independent of the MinD4 ATPase activity as mutation of the Walker A or Walker B motifs did not abrogate the oscillation although it altered its period. Patterning systems critically require an energy source to drive them away from equilibrium. In the *E. coli* Min system, energy is provided by MinD catalyzed ATP hydrolysis. As MinD4 continues to form an oscillating pattern when its ATPase function is inhibited, we infer that MinD4 ATPase is not driving patterning. This implies that MinD4 is most likely binding to another patterning protein system that is maintaining the oscillation in MinD4 concentration. Thus, *H. volcanii* must utilize the MinD4 ATPase for some other function which appears to affect directional movement.

**Figure 9:**
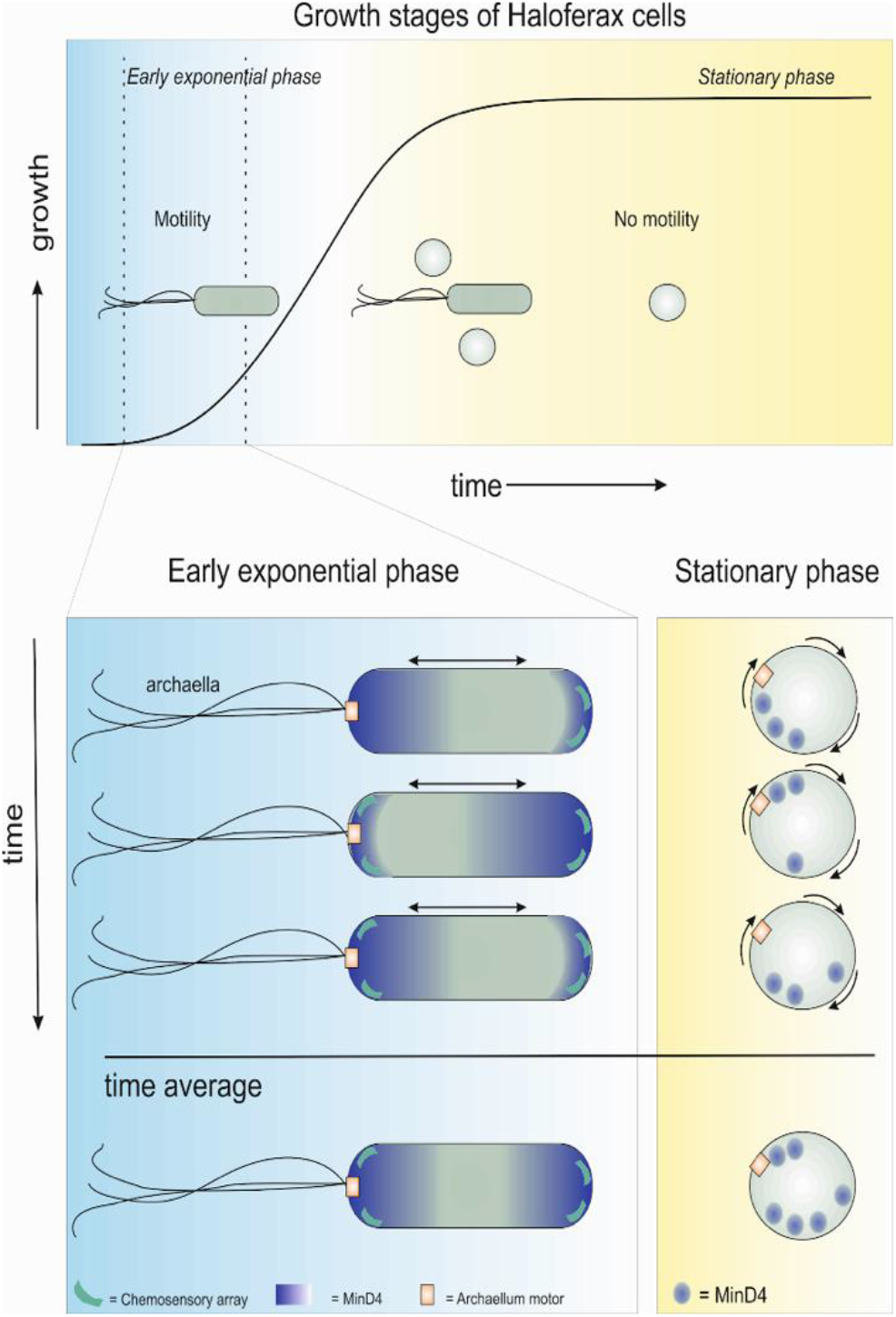
Schematic representation of MinD4 localization in *Haloferax* cells at different developmental stages. The top panel shows an idealized typical growth curve of *H. volcanii* cells, where cells progress from rod-shaped (blue), via a mixed population, to solely round cells. The motile phase is indicated between dotted lines. In the exponential growth phase, the MinD4 protein in found in patches at the cell pole, while a background distribution of MinD4 oscillates along the longitudinal cell axis. Archaella and chemosensory arrays are predominantly present at the cell poles of rod-shaped cells. In stationary phase the cells are round, and MinD4 is forming distinct foci that chase each other along the cell membrane.

MinD4 has an unusual long C-terminal domain, with the very C-terminal section homologous to the *E. coli* membrane targeting sequence (MTS)^28^. Deletion of the MTS or eMTS prevents the cellular oscillations in MinD4 concentration, thus the MTS plus a portion of the C-terminal domain is likely to be responsible for coupling MinD4 to the underlying oscillating system. In addition, MinD4_eMTS* and MinD4_MTS* are not distributed in polar patches, as is the case for wild type MinD4, but forms distinct foci. Thus, the MTS of MinD4 is important for oscillation, possibly by coupling MinD4 to another system or by correctly positioning it at the cell membrane. That MTS mutants were able to form foci but were not biologically functional (they displayed reduced motility) suggests, perhaps unsurprisingly, that coupling to the underlying patterning system is critical for MinD4 to function correctly.

In round (probably plate-like) cells, the MinD4 oscillation forms a rotating pattern, where the MinD4 concentration maximum rotates around the circumference of the cell (Fig 5F, 9). This is similar to patterning observed in the *E. coli* Min system when it is confined in artificial rectangular shapes^49^. Discoid *H. volcanii* cells were previously shown to be non-motile, and chemosensory arrays and archaella are not present in these cells (Fig 9) suggesting that the function of MinD4 in round cells is inconsequential^44^.

The archaeal MinD4 protein seems to combine traits from several bacterial ParA/MinD protein systems. It displays an oscillating distribution, as is the case for MinD proteins in gram-negative bacteria, and in addition forms polar patches, such as ParA homologs working in concert with a landmark protein for positioning at the cell pole (Fig 9). The oscillatory distribution of MinD4 might be the result of its interaction with another oscillating protein, while the formation of polar patches might rely on ATP-dependent binding to a cell pole organizing factor or landmark. Archaeal MinD4 is important for correct placement of motility and chemotaxis machinery (Fig 9), and this function is reminiscent of that of ParA homologs that do not oscillate and instead bind to landmark proteins at the cell poles.

The distribution pattern of chemosensory arrays in *H. volcanii* superficially resembles that of *E. coli*, although there are some apparent differences^44^. Several factors are suggested to play a role in the correct spatiotemporal positioning of chemosensory arrays of *E. coli*, such as: free diffusion and capture of receptors^56–58^, a preference for membrane curvature^59^,^60^, interactions with the Tol-Pal system^61^ and others. In *E. coli*, ParA/MinD seem not important for the positioning of chemosensory arrays^56^. As a fraction of the cell population of *ΔminD4 H. volcanii* still maintains correctly positioned archaella and chemosensory arrays, it might be possible that in *H. volcanii*, as in *E. coli*, a combination of multiple factors underlie their positioning pattern.

From the phenotype of the Δ*minD4* strain, the mechanism by which MinD4 affects formation of chemosensory arrays and archaella is not directly obvious. The ATP dependent formation of polar patches by MinD4 is important for the correct formation of chemosensory arrays and archaella, and thus for motility. Possibly the formation of polar patches reflects correct binding to a pole organizing factor. A clue as to the potential pole organizing factor to which MinD4 could bind, comes from previous cryo-EM studies. Archaella and chemosensory arrays in archaea, were found to be attached to a so-called ‘polar cap’, which is a large macromolecular sheet below the membrane at the cell poles, in which both archaella and chemosensory arrays seem to be anchored^62,63^ (Fig S8A). The proteins responsible for polar cap formation have not been identified yet.

We observed a decrease in the number of both cells with chemosensory arrays (from 80% to 40 %) and archaella (from 80 % to 20 %). Previously the correct positioning of both structures was shown to be independent from each other, and suggested to rely on an independent pole organizing factor^44^. Possibly, MinD4 is important for formation or positioning of the polar cap, and as such affects the formation of chemosensory arrays and archaella. The protein interaction network of MinD4 awaits further experimental characterization, and could help to identify a possible landmark or pole organizing factor.

Our first characterization of the biological function of archaeal MinD proteins shows similarities with that of different bacterial ParA/MinD homologs. Instead of being responsible for positioning the cell division ring, as is the case for bacterial MinDs, in the model euryarchaeal *Haloferax*, the MinD4 protein is involved in positioning of motility and chemotaxis machinery. This might be a peculiarity of the *Haloferacales*, or could indicate that all MinD encoding archaea use these proteins to position chemotaxis and motility machinery. In that case, euryarchaea might have developed other systems to correctly place the FtsZ ring.

## Acknowledgements

PN was supported by a grant from the Federal Ministry for Economic Affairs and Energy based on a decision of the German Bundestag (ZF 4653901AJ8). FD received support from the CRC746 funded by the DFG (German research foundation). MP received funds from the DFG on grant AL1206/4-3. TQ was supported by the DFG (411069969). ID and SI were supported by the Australian Research Council (FT160100010 and DP160101076).

## Author contributions

PN constructed all MinD deletion mutants and performed growth experiments and the analysis of CheW localization. SI performed all FtsZ localization studies, cell morphology analysis and MinD4 truncations. JW performed all oscillation analyses. FD constructed expression strains and oscillation analyses. MP performed motility assays, the analysis of FlaD localization and GFP Western Blot analysis. ID performed molecular phylogeny. TEFQ performed swimming microscopy and MFR electron microscopy. TQ, ID and SVA wrote the manuscript. All authors conceived the experiments and contributed to writing of the manuscript.

## Materials and Methods

### Identification of MinD superfamily proteins in archaea and phylogenetic analysis

To identify MinD superfamily (SIMIBI) genes, a search set of 14 homologs from well-characterized bacteria were aligned with MUSCLE ^64^ and used as the query set for searches of selected archaeal genomes by searching their respective proteome reference or UniProt databases using JackHMMER ^65^. Once the iterative JackHMMER searches for each species had converged, significant hits were selected that contained the deviant Walker A motif, (K/R)GGXG(K/R), allowing one conservative mismatch in the last 5 residues. Several short fragments were also removed. The species analyzed, and numbers of homologs identified in each, that conform to the above criteria, are given in Supplementary Table 1.

To identify the affiliation of each homolog with the subfamilies MinD, ParA, FlhG, Mrp/NBP35 or ArsA, the full set of homologs identified in each archaeon were aligned with Clustal-Omega ^66^ together with a representative set of homologs of each of the five families compiled from sets of both “top hits” and “most-diverse” representatives obtained from the Conserved Domain Database ^67^. The identification of homologs in specific archaeal species was then judged based on their position amongst the representative families in a Maximum likelihood phylogenetic tree, obtained using MEGA (ver. 7.0.26) ^68^, with 100 bootstrap replicates and using only those positions (columns) in the alignment that contained <50% gaps.

### Growth and genetic modification of H. volcanii

*Haloferax volcanii* was grown and genetically manipulated as described in^46,69^. Depending on the experiment, the cells were grown at 45 °C or 42 °C in either rich YPC medium with Bacto^™^ yeast extract, peptone (Oxoid, UK) and Bacto^™^ Casamino acids (BD Biosciences, UK or Oxoid) or in selective CA medium containing only Bacto^™^ casamino acids ^46,69^, occasionally supplemented with a trace elements solution as described ^43^ was also used where indicated (CAB medium). Plates were incubated at 45°C and liquid cultures were shaken at 120-200 rpm. For analysis with microscopy cells were diluted in CA medium with trace elements (CAB) and 0.2 mM tryptophan at 45 °C to maintain cells in exponential growth phase (0.350<OD_600_<0.550). These conditions were applied for coulter cytometry, high resolution fluorescent microscopy, fluorescent microscopy of MinD4 truncations and FtsZ1-GFP and analysis of cell shape by phase contract microscopy. For fluorescent microscopy of CheW and MinD4, overnight cultures were diluted in CA and CA with trace elements, respectively, and imaged the next day during different phases of exponential growth after one hour of induction with 1 mM tryptophan (from OD 0.01-0.5). Gene deletion and gene expression studies were generally carried out as described previously ^69^. Primers to create knockout plasmids, based on pTA131, are described in Table S3. To express fusion proteins, plasmids harbouring the *pyrE2* cassette were constructed (Table S2). These plasmids contained *mCherry* or *gfp* genes and in-frame restriction sites to enable the expression of N-terminal and C-terminal fluorescent fusion proteins under the control of the tryptophan promoter (Table S2).

### Coulter cytometry

The cell volume was analyzed with a Multisizer 4 Coulter Counter (Beckman Coulter, Brea, California, USA) equipped with a 30 nm aperture tube. Runs were completed in volumetric mode (100 μL), with the current set to 600 μA and gain set to 4. Samples from cultures maintained in exponential growth phase were diluted (1:100) in 18% Buffered Salt Water BSW (containing per liter 144 g NaCl, 21 g MgSO4.7H2O 429, 18 g MgCl2.6H2O, 4.2 g KCl,12 mM Tris-HCl (pH 7.4).

### Phase-contrast microscopy and cell shape analysis

Cells were placed on an agarose pad containing 1% agarose in 18% BSW. Images were acquired on a GE DV Elite microscope equipped with an Olympus 100X UPLSAPO/NA 1.4 objective and a pco.edge 5.5 sCMOS camera. Image analysis (segmentation and shape analysis) were performed with Fiji/MicrobeJ ^70,71^. Results are shown as a frequency distribution (calculated in Graph Pad Prism 8) grouped in bins of 0.05 intervals from 0 to 1 for the cell circularities and bins of 0.1 intervals from 0 to 12 for the cell areas.

### Fluorescence microscopy

Images were acquired on a GE DV Elite microscope equipped with an Olympus 100X UPLSAPO/NA 1.4 objective and a pco.edge 5.5 sCMOS camera and a GFP/FITC filter (ex:464-492 nm em:500-523 nm) for FtsZ1 and MinD4 truncations, and on a Zeiss Axio Observer 2.1 Microscope, (ex: 450-490 nm em: 500-550 nm filter from Chroma^®^), equipped with a heated XL-5 2000 Incubator running VisiVlEW℗ software for MinD4 and CheW. Exposure time of MinD4 and CheW was 2 sec. Exposure time of GFP-MinD4ΔMTS and GFP-MinD4ΔeMTS was 0.6 sec at 50% intensity.

### Image analysis

Fiji and MicrobeJ ^70^,^71^were used to calculate the relative median intensity of FtsZ1-GFP in the diverse knock out mutants, the relative cellular positioning of GFP-MinD4 and mutants thereof, and the localization and mobility of GFP-CheW. FtsZ1-GFP Fluorescence channels were first treated with a background subtraction filter (ballsize=8) in Fiji before image analysis with MicrobeJ. The relative median intensities of FtsZ1-GFP as a function of the relative long cell axis (represented from 0 to 1) were exported to Graph Pad Prism 8.0 then normalized and the % of total was calculated before plotting. GFP Fluorescence channels of the MinD4 truncations were treated with a white top hat filter from the Fiji’s MorpholibJ plugin (ballsize=8) before image analysis with MicrobeJ. The relative localization of each detected focus of MinD and mutants thereof was plotted as a function of the relative cell shape using MicrobeJ. The long axis was defined as the line of greatest distance across the cell in which it splits the cell into two equal areas. Polarity of cells were not defined. The foci positions were then mapped relative to this axis in an idealized oblong-shaped cell.The percentage of cells with foci and the number of foci per cell was calculated with the MicrobeJ Statistics tool and data exported for plotting in GraphPad Prism 8. For each experiment, the cells were grown and observed at least on three independent occasions, resulting in the analysis of at least several hundred cells.

### Time lapse microscopy

For the live imaging of *H. volcanii* cells, 0.3 % (w/v) agarose pads made of CAB with 1 mM tryptophan were poured in round DF 0.17 mm microscopy dishes (Bioptechs). The agar pad was removed after drying and the cells were placed under the agar pad, after which the lid was placed on the microscopy dish. The microscope chamber was heated at 45 ° C. Images in the PH3 and GFP modes were captured at 100 × magnification every 3 minutes for 1 hour, or every 30 min for 16 hours.

Time lapse movies for quantitative analysis of localization were collected using epifluorescence on a custom built microscope based around an ASI-RAMM frame (Applied Scientific Instrumentation) with a Nikon 100 × CFI Apochromat TIRF (1.49 NA) oil immersion objective. Lasers were incorporated using the NicoLase system^72^. Images were captured on an Andor iXon 888 EMCCD cameras (Andor Technology Ltd). 300 mm tube lenses were used to give a field of view of 88.68 μm × 88.68 μm. Timelapse movies were acquired at 45°C using an Okolab stage top environmental chamber. Cells were prepared as per cell shape measurements with samples held in a Chamlide Chamber. Movies were acquired for half an hour at one frame per 10 seconds using 100 ms exposures with 30 mW 488 nm laser illumination.

Calculation of parameters of oscillation: Cell shape outlines and their respective lowest harmonic modes were detected using software available publicly on Github at: https://github.com/lilbutsa/Archaea_Division_Analysis as described in^45^. The magnitude of oscillations were calculated by taking the inner product of the lowest harmonic mode of each cell shape with their respective fluorescent signal for each frame^73^. Oscillations periods were calculating by measuring twice the time between zero crossing points in the patterning magnitude after a Gaussian filter of width 5 frames was applied to filter noise. Amplitudes were taken as the maximum absolute magnitude of oscillation between each crossing time. The final amplitude and magnitude of each cells patterning was found by taking the mean of the individually measured oscillations.

### Motility analysis on semi-solid agar plates

Semi-solid agar plates were made of 0.33% agar in CA medium with 1 mM tryptophan. Over-night cultures with an OD of ~0.5 were stabbed on the plates and plates were incubated for 4 days at 45 °C. All strains of which the motility was compared with each other, were spotted on the same plate. Of each strain, at least 3 technical and 3 biological replicates were performed. After 4 days, the diameter of the motility ring was measured. The diameter of the reference strain was set to 100% and that of the other strains was measured relative to that. All values of all independent experiments were subjected to a two tailed unpaired T-test, in comparison with the wt strain.

### Swimming analysis by live microscopy

Cell motility was analyzed by live microscopy as described in ^46^. Briefly, precultures of cells were grown over night in CA medium with uracil at 45 °C. The next day, cells were diluted to a theoretical OD of 0.005 in 20 mL CA with uracil and grown for ~16 hours until the OD was ~0.05. Cells were transferred to a DF 0.17 mm microscopy dish (Bioptechs) and observed at 40x magnification in the PH2 mode with a Zeiss Axio Observer 2.1 Microscope running with VisiVIEW℗software at 45 °C. Time-lapse movies were made during 15 seconds with 20 frames per second.

### Western blotting

Similar cultures as analyzed by fluorescence microscopy were harvested to test for the stability and correct expression of the GFP fusion proteins. Cell cultures were harvested at 5000 rpm for 20 min, and the pellet was re-suspended in 1x SDS buffer (250 mM Tris pH 6.8, 10% SDS(w/v), 10% Dithiothreitol (DTT) (w/v), 50% glycerol (v/v), 50 μg/mL) concentrated in PBS (Phosphate Buffered Saline) to a theoretical O.D of 22. 10μL of the sample was loaded and run on a 11% SDS-PAGE. After gel electrophoresis, the proteins were transferred to a PVDF membrane using semidry blotting in blotting buffer (5 mM Tris, 40 mM glycine, 20 (v/v), 0.0375% (v/v)). The membrane was blocked with 0.1 % I-Block^™^ (Applied Biosystems, California USA) for 1 hour at room temperature. Next, the membrane was incubated in primary antibody, α GFP (Sigma-aldrich, California USA; 1:5000 dilution) for 3-4 hours at 4°C. After incubation, the membrane was washed 3 times with PBST. The secondary antibody, α-rabbit coupled to Horse Radish Peroxidase (from goat) (ThermoFischer Scientific, Masasachusetts USA; 1:10,000 dilution) was added and incubated for 1 hour at 4°C. The membrane was washed 3 times with PBST after the incubation period, and prepared for development. Invitrogen ^™^ I-Bright^™^ (ThermosFischer Scientific) scanner was used for image development.

### Electron microscopy

Cells for electron microscopy were grown as described for ‘swimming analysis by live microscopy’. Cells were centrifuged at 2000g for 10 min and concentrated to a theoretical OD of 20 in CA medium and fixed in 2.5% GA and 1 % FA. Cells were spotted on glow discharged carbon-coated grids containing Formvar films, by placing 5 μl of the cell-suspensions on the grid for 30 sec. Samples were washed three times with distilled H2O and negatively stained for 30 sec with 2% (wt/vol) uranyl acetate. Cells were imaged using a Philips CM10 transmission EM coupled to a Gatan 792 BioScan camera and the Gatan DigitalMicrograph software.

### Primers, plasmids, strains

Primers, plasmids and strains used in this study are described in Table S2-S4.

