## Supplementary Material for "An oscillating MinD protein determines the cellular positioning of the motility machinery in archaea"

# to

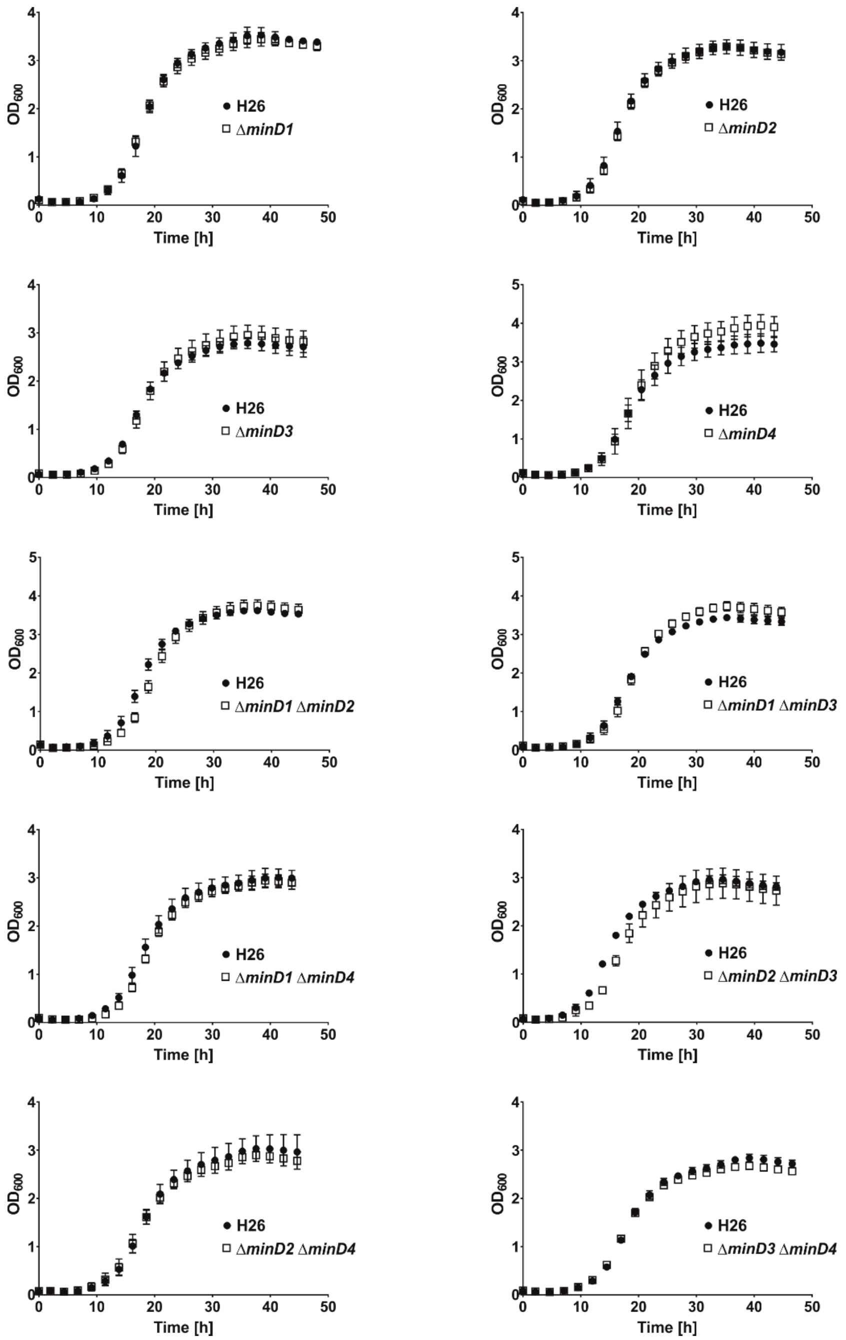

**Figure S1 MinD homologs do not influence growth of *H. volcanii***. The $OD_{600}$ of at least three biological replicates of wild type (H26) and several Δ*minD* strains in liquid CAB medium was monitored for 45 hours. Error bars, SD.

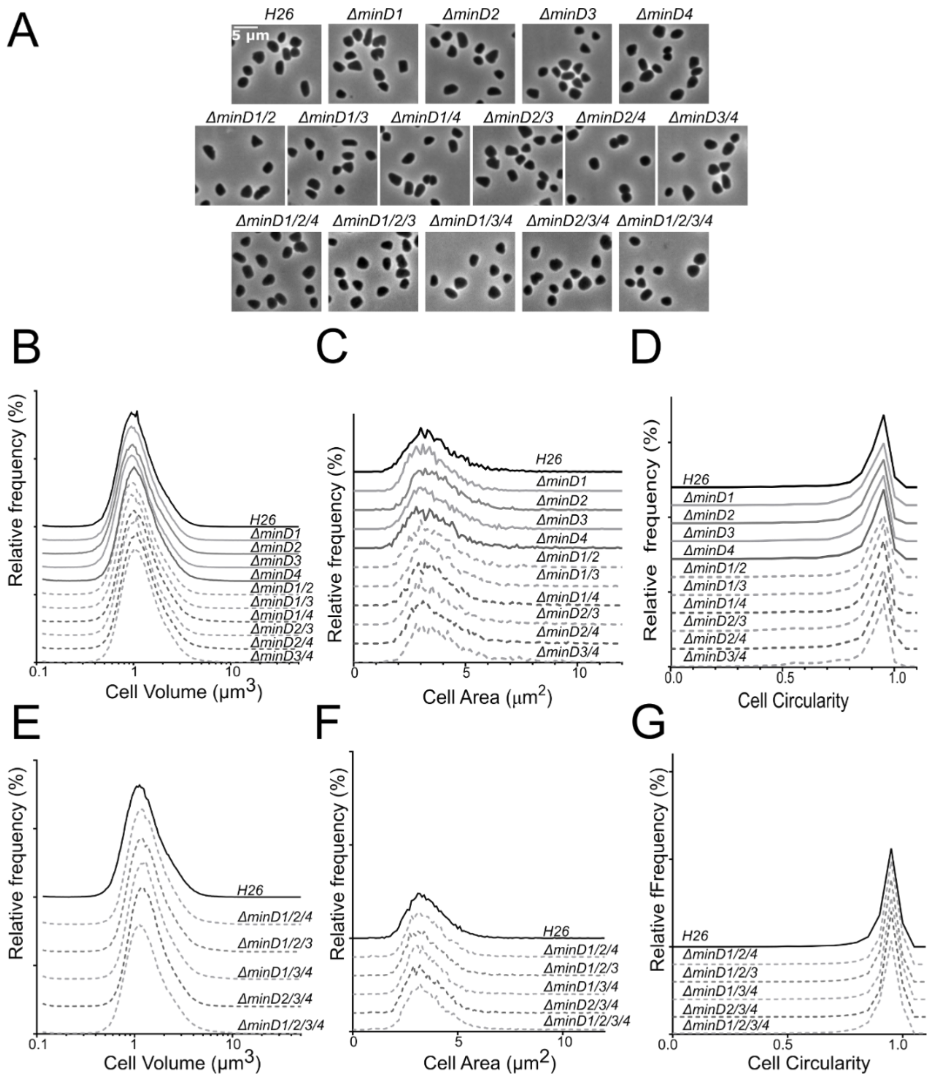

**Figure S2.** **Cell shape and division are not affected by deletion of MinD proteins in *H. volcanii***

(A) Phase contrast images of different *H. volcanii* minD mutant strains. (B) and ( E) Relative frequency distributions of cell volume measured by Coulter cytometry. (C) and (F) Relative frequency distributions by automated image analysis of cell area. (D) and (G) Relative frequency distributions by automated image analysis of cell circularity. Cells sampling are performed on cultures in logarithmic growth phase in CAB medium. All results are the average of at least two representative independent experiments. Sample sizes for C and E : $n_{H26}$ =3235, $n_{\Delta minD1}$=3044, $n_{\Delta minD2}$= 3113, $n_{\Delta minD3}$=2889, $n_{\Delta minD4}$=2568, $n_{\Delta minD1/2}$=2582, $n_{\Delta minD1/3=3633}$, $n_{\Delta minD1/4=2689,}$ $n_{\Delta minD2/3=3839}$, $n_{\Delta minD2/4=3459}$, $n_{\Delta minD3/4=2590}$ ; F and G : $n_{H26}$ =5730 , $n_{\Delta minD1/2/4}$=8248, $n_{\Delta minD2/3/4}$=4583, $n_{\Delta minD1/3/4}$=5825, $n_{\Delta minD1/2/3}$=4113, $n_{\Delta minD1/2/3/4}$=10930.

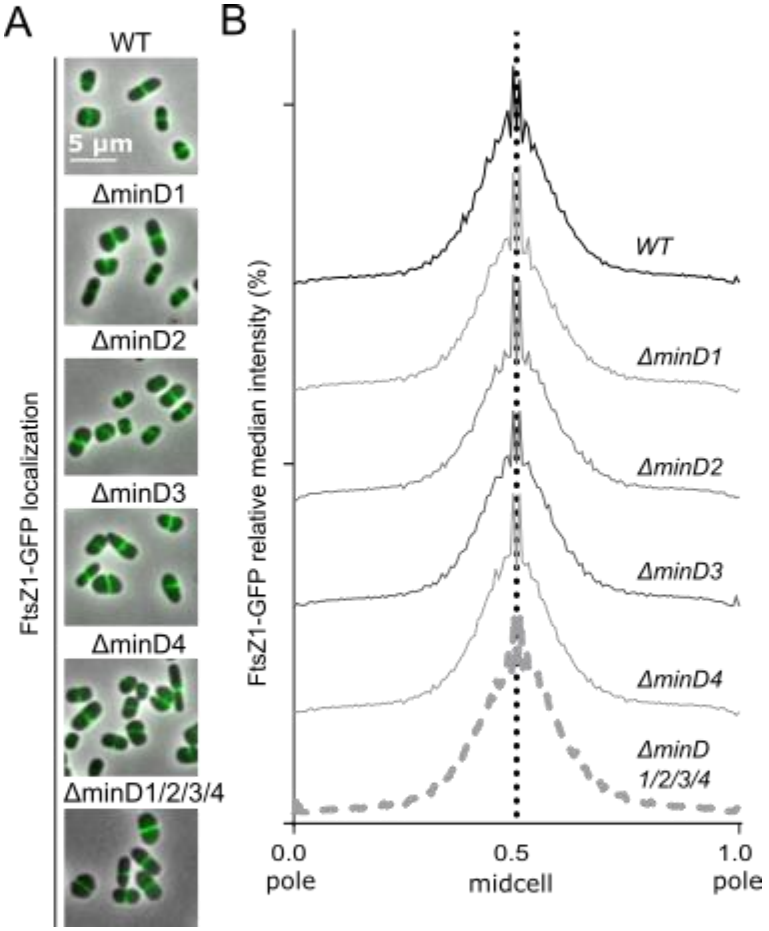

**Figure S3 MinD homologs of *H. volcanii* do not affect positioning of FtsZ1.** (A) FtsZ1-GFP localization in single and quadruple *minD* mutants during exponential growth phase. Figures show an overlay of phase contrast and GFP fluorescence channels. (B) Representation of the relative median intensity of the FtsZ1-GFP signal along the relative cell length measured by automated image analysis. Cells sampling are performed on cultures in logarithmic growth phase in CAB medium. Sample sizes: $n_{WT}$=1252, $n_{\Delta minD1}$=1485, $n_{\Delta minD2}$=3588, $n_{\Delta minD3}$=3529, $n_{\Delta minD4}$=3950; $n_{\Delta minD1/2/3/4}$=2747.

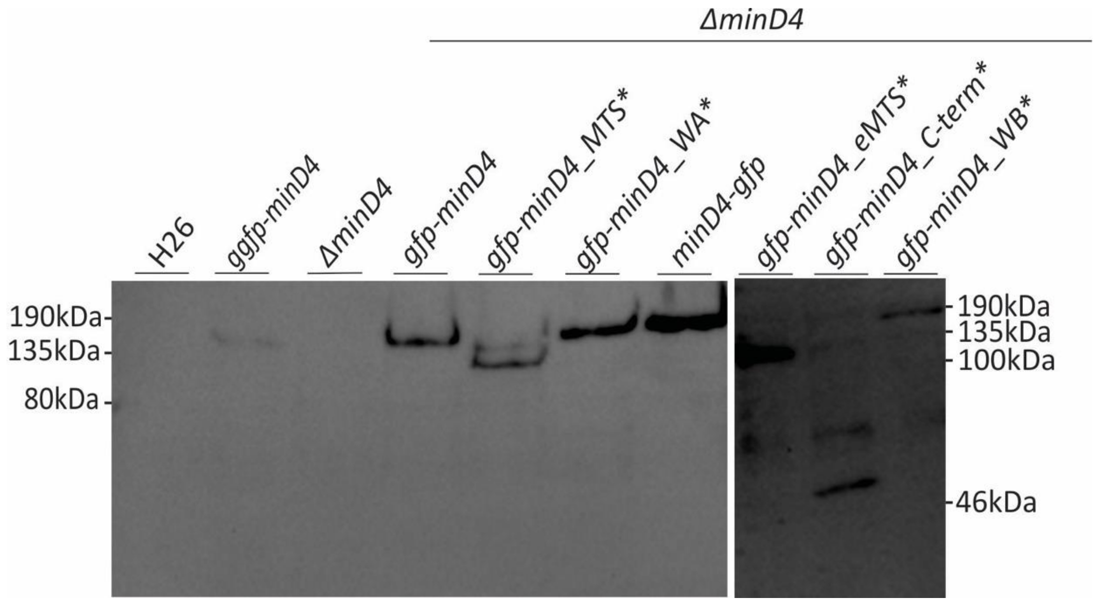

**Fig S4 Expression of GFP-MinD4 fusion proteins.** Western blot analysis with a primary α-GFP antibody from rabbit and a secondary HRP coupled antibody on total cell lysates of *H. volcanii* cells in exponential growth phase, expressing different fusion proteins. Cells were grown under similar conditions as used for fluorescent microscopy. The position of marker proteins of known size is indicated on the side of the blot.

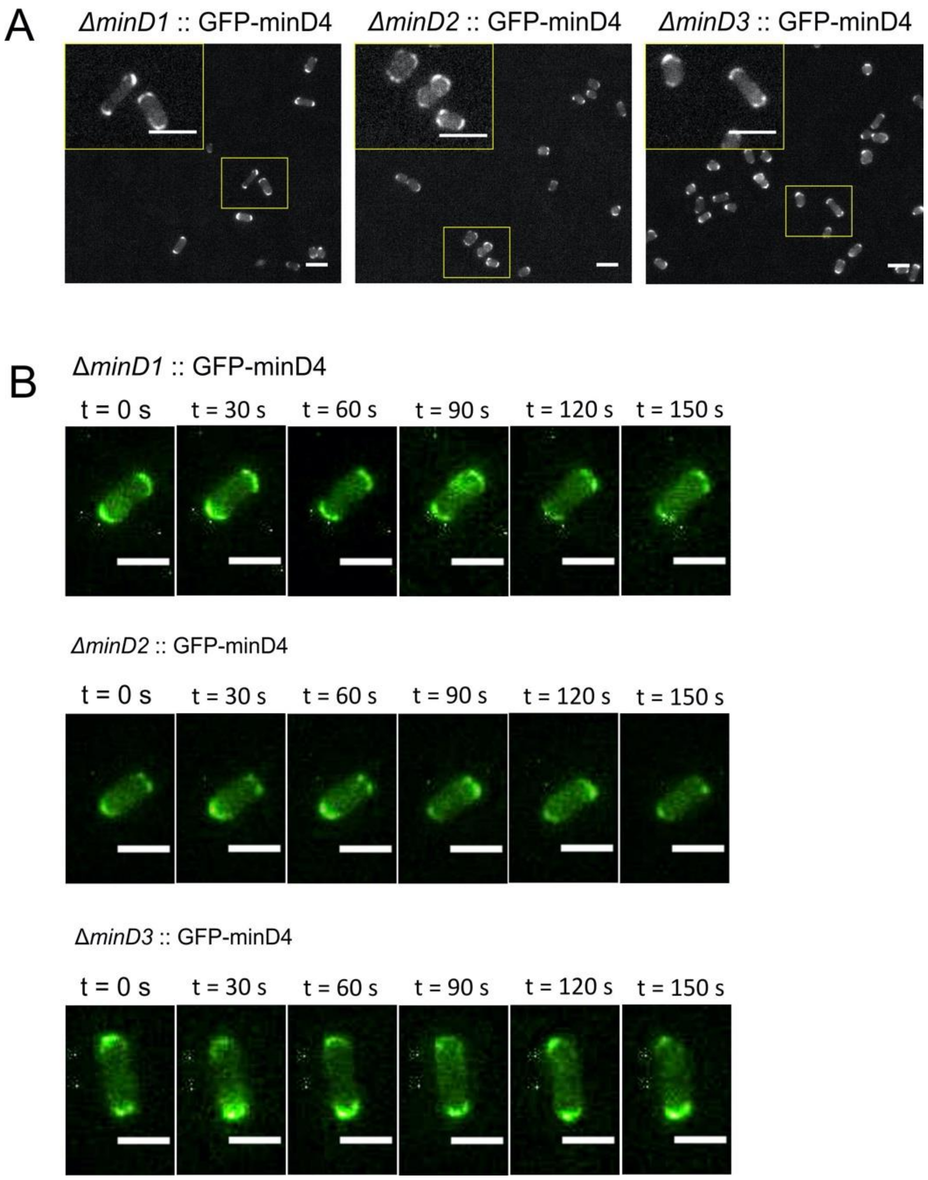

**Fig S5. MinD4 localization and oscillation are not dependent on the presence of other MinD homologs in *H. volcanii*.** (A) Localization pattern of pGFP-MinD4 in the *H. volcanii* Δ*minD1*, Δ*minD2*, Δ*minD3* strains. Scale bar 4 µm (B) Still images of time lapse movies of representative cells of each strain expressing pGFP-MinD4. Scale bar, 4 µm

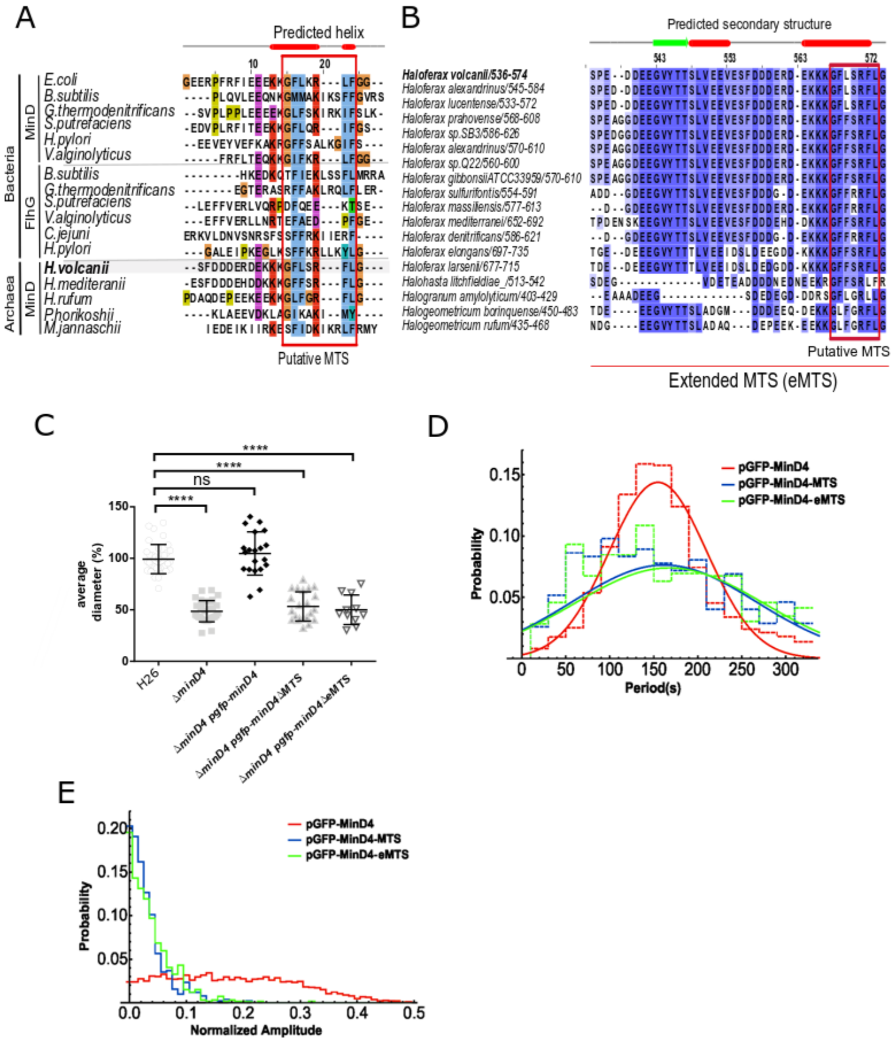

**Figure S6. MTS domain conservation in minD4 homologs and its importance for localization and oscillation**

(A) Amino acid sequence alignment of MinD homologs in archaea and bacteria showing the predicted conserved MTS in the red box. (B) Amino acid sequence alignment of the extended MTS (eMTS) region in the C-terminal region of the MinD4 homologs of different haloarchaea. (C) Average diameter of motility rings, measured relative to the wild type, from different *H. volcanii* strains from >3 independent experiments including 4 biological replicates each. Middle black line indicates mean, lower and upper lines the standard deviation. (D) The distribution of periods of oscillations observed for each variant of MinD4. (E) The relative amplitude distribution for each variant of MinD4 given by the amplitude of oscillations divided by the total fluorescence within the cell. MinD4-MTS (AA 1-560), MinD4-eMTS (AA 1-535).

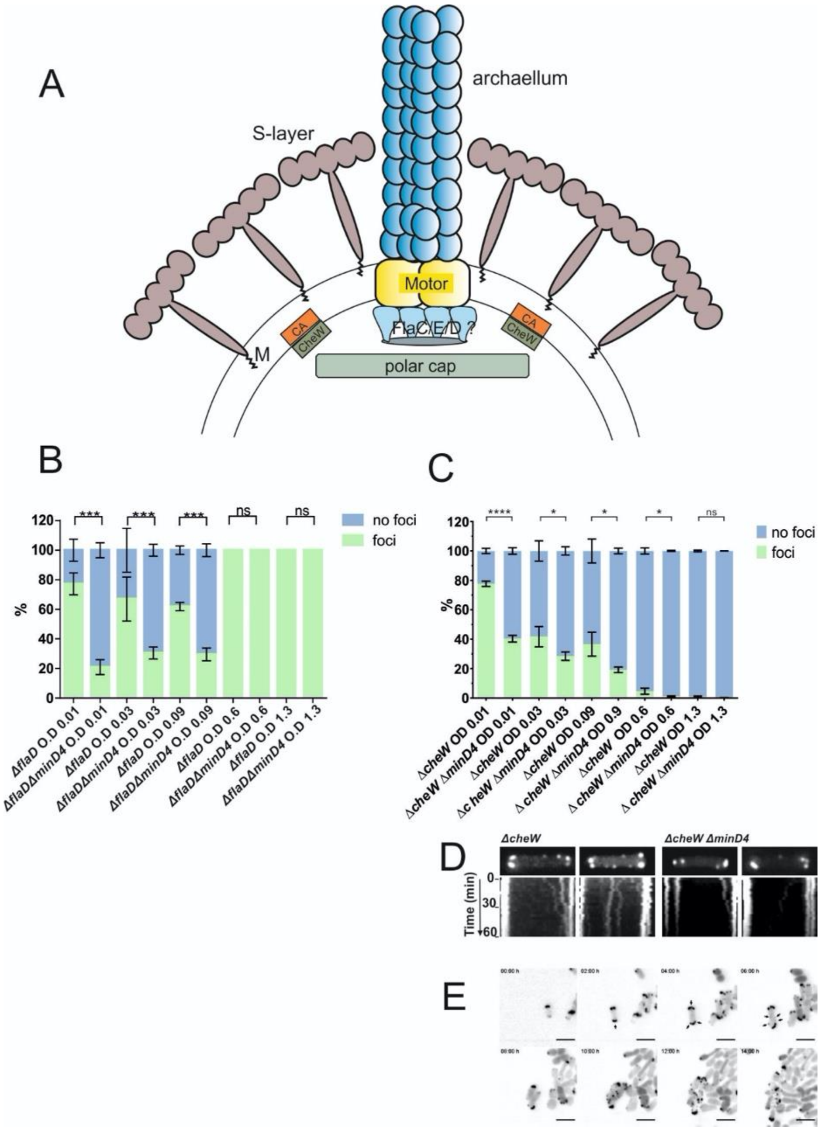

**Figure S7: The cellular positioning of FlaD and CheW is affected by MinD4**

(A) Model of the assembled archaellum at the cell pole of *H. volcanii*. CA, chemosensory arrays; M, membrane. (B) Presence of FlaD-GFP foci in the Δ*flaD* and the Δ*flaD*Δ*minD4* strain at different growth stages (OD0.01- 1.3). (C) Presence of of GFP-CheW foci in the Δ*cheW* and the Δ*cheW*Δ*minD4* strain at different growth stages (OD0.01- 1.3). (D) Dynamics of GFP-CheW in the ΔcheW and the ΔcheWΔminD4 strain. Upper panel: representative cells with GFP-CheW localization. Lower panel: kymograph of representative cells shown in the upper panel. Mobility of GFP-CheW followed during 1 hour. (E) Time lapse movie of GFP-CheW in the ΔcheWΔminD4 strain during cell division over 16 hours. Scale bar, 4 µm.

Supplemental Movies

Movie S1: Time-lapse of rod-shaped gGFP-MinD4 *H. volcanii* cell

Movie S2: Time-lapse of a rod-shaped Δ*minD4 H. volcanii* cell expressing pGFP-MinD4

Movie S3: Time-lapse of a round Δ*minD4 H. volcanii* cell expressing pGFP-MinD4

Movie S4: Time-lapse of a Δ*minD4 H. volcanii* cell expressing pGFP-MinD4_WA*

Movie S5: Time-lapse of a Δ*minD4 H. volcanii* cell expressing pGFP-MinD4_WB*

Movie S6: Time-lapse of a Δ*minD4 H. volcanii* cell expressing pGFP-MinD4_ΔMTS

Movie S7: Time-lapse of a Δ*minD4 H. volcanii* cell expressing pGFP-MinD4_ ΔeMTS

Movie S8: Time-lapse movie of 15 sec showing swimming of Δ*pilB3 H. volcanii* cells in liquid CA medium

Movie S9: Time-lapse movie of 15 sec showing swimming of Δ*pilB3* Δ*minD4 H. volcanii* cells in liquid CA medium

Movie S10: Time-lapse movie of 1 hour showing mobility of chemosensory clusters in Δ*cheW* Δ*minD4 H. volcanii* cells expressing pGFP-CheW

**Table S1.** Number of archaeal homologs identified with a bacterial ParA/MinD/FlhG search alignment*

| *Taxon represented* | *Example species/isolate* | *Abbreviation* | *Homologs (Dev. Walker A)* |
|---|---|---|---|
| Euryarchaeota | | | |
| Archaeoglobi | Archaeoglobus fulgidus DSM 4304 | ARCFU | 8 |
| | Ferroglobus placidus DSM 10642 | FERPA | 6 |
| | Geoglobus acetivorans (taxid: 565033) | GEOAC | 5 |
| Methanoliparia | Euryarchaeota archaeon NM1a (taxid:2491083) | EANM1 | 7 |
| Thermoplasmata | Thermoplasma acidophilum DSM 1728 | THEAC | 2 |
| | Methanomassiliicoccus luminyensis B10 | METLB | 12 |
| Hadesarchaea | Hadesarchaea archaeon DG-33 | HADES | 4 |
| Methanobacteria | Methanobrevibacter ruminantium M1 | METRM | 5 |
| | Methanobacterium lacus (taxid:877455) | METLA | 12 |
| | Methanosphaera stadtmanae DSM 3091 | METST | 4 |
| | Methanothermobacter thermautotrophicus str. Delta H | METTH | 10 |
| | Methanothermus fervidus DSM 2088 | METFV | 7 |
| Methanococci | Methanocaldococcus jannaschii DSM 2661 | METJA | 12 |
| | Methanococcus maripaludis S2 | METMP | 10 |
| Methanonatronarchaeia | Methanonatronarchaeum thermophilum | MENAT | 1 |
| Methanopyri | Methanopyrus kandleri AV19 | METKA | 8 |
| Nanohaloarchaeota | Candidatus Nanosalina sp. J07AB43 | NANS0 | 4 |
| | Candidatus Haloredivivus sp. G17 | HALSG | 1 |
| Halobacteria | Haloarcula japonica DSM 6131 | HALJA | 1 |
| | Natronomonas pharaonis DSM 2160 | NATPD | 6 |
| | Halobacterium salinarum NRC-1 | HALSA | 10 |
| | Halococcus saccharolyticus DSM 5350 | HALSC | 3 |
| | Natronoarchaeum philippinense | NATPH | 12 |
| | Haloferax volcanii DS2 | HALVD | 11 |
| | Haloquadratum walsbyi DSM16790 | HALWD | 10 |
| | Halorubrum lacusprofundi ACAM 34 | HALLT | 16 |
| | Natrialba magadii ATCC 43099 | NATMM | 8 |
| Methanomicrobia | Methanocella arvoryzae MRE50 | METAR | 9 |
| | Methanoculleus marisnigri JR1 | METMJ | 8 |

| | | | |
|---|---|---|---|
| | Methanophagales archaeon ANME-1-THS | ANME1 | 8 |
| | Methanosarcina acetivorans C2A | METAC | 14 |
| Theionarchaea | Theionarchaea archaeon DG-70 | THEIO | 7 |
| Thermococci | Pyrococcus furiosus DSM 3638 | PYRFU | 6 |
| | Thermococcus kodakarensis KOD1 | THEKO | 9 |
| | Palaeococcus pacificus DY20341 | PALPA | 10 |
| ***ASGARD GROUP:*** | | | |
| Heimdallarchaeota | Candidatus Heimdallarchaeota archaeon | HEIMD | 1 |
| Lokiarchaeota | Lokiarchaeum sp. GC14_75 | LOKSG | 7 |
| Odinarchaeota | Candidatus Odinarchaeota archaeon LCB_4 | ODINA | 2 |
| Thorarchaeota | Candidatus Thorarchaeota archaeon | THOAR | 9 |
| ***DPANN GROUP:*** | | | |
| Aenigmarchaeota | Candidatus Aenigmarchaeota archaeon CG1_02_38_14 | AENIG | 1 |
| Diapherotrites | Candidatus Diapherotrites archaeon | DIAPH | 2 |
| Micrarchaeota | Candidatus Micrarchaeum acidiphilum ARMAN-2 | MICA2 | 2 |
| Pacearchaeota | Candidatus Pacearchaeota archaeon CG1_02_30_18 | PACEA | 2 |
| Parvarchaeota | Candidatus Parvarchaeum acidiphilum ARMAN-4 | PARA4 | 2 |
| Woesearchaeota | Candidatus Woesearchaeota archaeon CG1_02_33_12 | WOESE | 2 |
| Nanoarchaeota | Nanoarchaeum equitans Kin4-M | NANEQ | 1 |
| ***TACK GROUP:*** | | | |
| Bathyarchaeota | Bathyarchaeota archaeon B23 | BATHA | 2 |
| Geothermarchaeota | Candidatus Geothermarchaeota archaeon ex4572_27 | GEOAR | 10 |
| Korarchaeota | Candidatus Korarchaeum cryptofilum OPF8 | KORCO | 4 |
| Marsarchaeota | Candidatus Marsarchaeota G1 archaeon BE_D | MARSA | 1 |
| Verstraetearcheota | Candidatus Methanosuratus sp. (taxid: 2495426) | VERST | 3 |
| Crenarchaeota | | | |
| Acidilobales | Acidilobus saccharovorans 345-15 | ACIS3 | 1 |
| Desulfurococcales | Aeropyrum pernix K1 | AERPE | 1 |
| Fervidicoccales | Fervidicoccus fontis Kam940 | FERFK | 7 |
| Sulfolobales | Sulfolobus acidocaldarius DSM 639 | SULAC | 3 |
| Thermoproteales | Pyrobaculum aerophilum str. IM2 | PYRAE | 2 |
| Thaumarchaeota | | | |
| Cenarchaeales | Cenarchaeum symbiosum A | CENSY | 1 |
| Nitrosopumilales | Nitrosopumilus maritimus SCM1 | NITMS | 2 |
| | Nitrosopumilales archaeon CG_4_10_14_0_8_um_filter_34_8 | NITAR | 1 |

| | | | |
|---|---|---|---|
| Nitrososphaeria | Candidatus Nitrososphaera evergladensis SR1 | NITES | 4 |

* Grey text indicates taxa in which genome data might be incomplete (*i.e.* in contig or scaffold form), or the taxon or species is *Candidatus* status.

Table S2 Strains used in this study

Table S2 Strains used in this study

| **Strain Name** | **Genotype** | **Reference** |
|---|---|---|
| ***H. volcanii*** | | |
| H26 | Δ*pyrE2* | [1] |
| HTQ 63 | Δ*pyrE2* Δ*cheW* | [2] |
| HTQ200 | Δ*pyrE2* Δ*minD1* | This |
| HTQ214 | Δ*pyrE2* Δ*minD3* | This study |
| HTQ218 | Δ*pyrE2* Δ*minD4* | This study |
| HTQ228 | Δ*pyrE2* Δ*minD2* | This study |
| HTQ229 | Δ*pyrE2* Δ*minD1* Δ*minD3* | This study |
| HTQ240 | Δ*pyrE2* Δ*minD1* Δ*minD4* | This study |
| HTQ241 | Δ*pyrE2* Δ*minD2* Δ*minD4* | This study |
| HTQ242 | Δ*pyrE2* Δ*minD3* Δ*minD4* | This study |
| HTQ243 | Δ*pyrE2* *gfp_minD4* | This study |
| HTQ244 | Δ*pyrE2* Δ*minD4* Δ*flaD1* | This study |
| HTQ245 | Δ*pyrE2* Δ*minD2* Δ*minD3* | This study |
| HTQ246 | Δ*pyrE2* Δ*minD1* Δ*minD2* | This study |
| HTQ247 | Δ*pyrE2* Δ*pilB3* | This study |
| HTQ249 | Δ*pyrE2* Δ*minD4* Δ*pilB3* | This study |
| HTQ257 | Δ*pyrE2* Δ*minD4* Δ*cheW* | This study |
| HTQ 258 | Δ*pyrE2* Δ*minD1* Δ*minD2* Δ*minD4* | This study |
| HTQ 259 | Δ*pyrE2* Δ*minD2* Δ*minD3* Δ*minD4* | This study |
| HTQ 260 | Δ*pyrE2* Δ*minD1* Δ*minD3* Δ*minD4* | This study |
| HTQ 261 | Δ*pyrE2* Δ*minD1* Δ*minD2* Δ*minD3* | This study |
| HTQ262 | Δ*pyrE2* Δ *minD1* Δ*minD2* Δ*minD3* Δ*minD4* | This study |
| ***E. coli*** | | |
| 10-beta Competent Cells | Δ*(ara-leu) 7697 araD139 fhuA* Δ*lacX74 galK16* | New England Biolabs |
| *dam*$^-$/*dcm*$^-$ competent | *ara-14 leuB6 fhuA31 lacY1 tsx78 glnV44 galK2* | New England Biolabs |

Table S3 Plasmids used in this study

| **Plasmids** | **Description** | **Primers used** | **Enzymes used** | **Source/reference** |
|---|---|---|---|---|
| pTA131 | Integrative plasmid with a *pyrE2* selection marker for knock-outs in *H. volcanii* (Amp$^r$) | - | - | [3] |
| pTA1392 | Overexpression plasmid for *H. volcanii* under the control of a tryptophan inducible promotor. Contains *pyrE2* selection marker (Amp$^r$) | - | - | [4] |
| pIDJL40 | Plasmid to express proteins with a C-terminal gfp phusion under the control of a tryptophan inducible promotor. Contains *pyrE2* selection marker (Amp$^r$) | - | - | [5] |
| pSVA1839 | Integrative plasmid with a *pyrE2* selection marker to knock-out *hvo_0225* (*minD1*) (Amp$^r$) | 6911, 6912: 6913, 6914 | KpnI, NdeI : NdeI, XbaI | This study |
| pSVA1840 | Integrative plasmid with a *pyrE2* selection marker to knock-out *hvo_0322* (*minD4*) (Amp$^r$) | 6915, 6916: 6917, 6918 | KpnI, NdeI : NdeI, XbaI | This study |
| pSVA1841 | Integrative plasmid with a *pyrE2* selection marker to knock-out *hvo_0595* (*minD2*) (Amp$^r$) | 6919, 6620: 6921, 6922 | KpnI, NdeI : NdeI, XbaI | This study |
| pSVA1842 | Integrative plasmid with a *pyrE2* selection marker to knock-out *hvo_1634* (*minD3*) (Amp$^r$) | 6923, 6924: 6925, 6926 | KpnI, NdeI : NdeI, XbaI | This study |
| pSVA3916 | Plasmid to express *minD4* with a C-terminal *gfp*-tag (Amp$^r$) | 6562, 6563 | NdeI, BamHI | This study |
| pSVA3919 | Plasmid to express *flaD1* with a C-terminal gfp-tag (Amp$^r$) | - | - | [2] |
| pSVA3922 | Plasmid to express proteins with an N-terminal GFP phusion under the control of a tryptophan inducible promotor. Contains *pyrE2* selection marker (Amp$^r$) | 6597, 6598 | NdeI, NheI | This study |

| | | | | |
|---|---|---|---|---|
| pSVA3924 | Plasmid to express *minD4* with a N-terminal *gfp*-tag (Amp$^r$) | 6599, 8000 | NheI, BamHI | This study |
| pSVA3950 | Plasmid to express truncated *minD4* (1-550 AA) with a N-terminal *gfp*-tag (Amp$^r$) | 6599, 8081 | NheI, BamHI | This study |
| pSVA3978 | Plasmid to express *minD4* (K16A) with a N-terminal *gfp*-tag (Amp$^r$) | 8826, 8827 | - | This study |
| pSVA3979 | Plasmid to express *minD4* (D117A) with a N-terminal *gfp*-tag (Amp$^r$) | 8828, 9728 | - | This study |
| pSVA3987 | Integrative plasmid with a *pyrE2* selection marker to integrate a genomic *gfp*-tag at the 5' start of *hvo_0322* (*minD4*) (Amp$^r$) | 8847, 8848: 8849, 8850 | - | This study |
| pSVA3999 | Integrative plasmid with a *pyrE2* selection marker to knock-out *hvo_1034* (*pilB3*) (Amp$^r$) | 8871, 8872: 8873, 8874 | KpnI, BamHI: BamHI, XbaI | This study |
| pSVA5031 | Expression plasmid for CheW-GFP | - | - | [2] |
| pHVID111 | GFP-MTS (MinD4 $_{R561-G574}$) expression plasmid | SI49, SI50 | NheI, BamHI | This study |
| pHVID112 | GFP-eMTS (MinD4 $_{S536-G574}$) expression plasmid | SI77, SI78 | NheI, BamHI | This study |
| pHVID113 | GFP-Cterm (MinD4 $_{T233-G574}$) expression plasmid | SI75, SI76 | NheI, BamHI | This study |

Table S4 Primers used in this study

| **Primer Number/Name** | **Sequence 5'→3'** | **Description** |
|---|---|---|
| 6562 | GTTCTACATATGGCCCGGGTGTACGC | Forward primer for the amplification of *hvo_0322* with NdeI restriction site |
| 6563 | GTAGGATCCGCCGAGGAAGCGACTGAG | Reverse primer for the amplification of *hvo_0322* with BamHI restriction site |
| 6597 | GTTCTACATATGAGTAAAGGAGAAGAAC | Forward primer for the amplification of *gfp* with NdeI restriction site |
| 6598 | GTAGCTAGCTTTGTATAGTTCATCCATGC | Reverse primer for the amplification of *gfp* with NheI restriction site |
| 6599 | AGTGCTAGCGCCCGGGTGTACGCAGTTG | Forward primer for the amplification of *hvo_0322* with NheI restriction site |

| | | |
|---|---|---|
| 8000 | CTAGGATCCAGACCTTCCCGTTTAGCC | Reverse primer for the amplification of *hvo_0322* with BamHI restriction site |
| 6911 | GATAGGTACCCGGTTCCCGTTTCTGCTGAC | Forward primer for the amplification of *minD1* (*hvo_0225*) up-stream flanking region with a KpnI restriction site |
| 6912 | GTATCATATGTCACGCGAGCCTCCGGAG | Reverse primer for the amplification of *minD1* (*hvo_0225*) up-stream flanking region with a NdeI restriction site |
| 6913 | GTCACATATGAAGCGTCGTCTCGTCTTCG | Forward primer for the amplification of *minD1* (*hvo_0225*) down-stream flanking region with a NdeI restriction site |
| 6914 | GTACTCTAGACGCCCAGAACGAAGACCTCC | Reverse primer for the amplification of *minD1* (*hvo_0225*) down-stream flanking region with a XbaI restriction site |
| 6915 | CTAGGGTACCGCGGGCTTTCGAGTACCTTG | Forward primer for the amplification of *minD4* (*hvo_0322*) up-stream flanking region with a KpnI restriction site |
| 6916 | CATGCATATGAATTCCAACGTCGAAGTCCAC | Reverse primer for the amplification of *minD4* (*hvo_0322*) up-stream flanking region with a NdeI restriction site |
| 6917 | GGTTCATATGTAAACGGGAAGGTCTTTTACTC | Forward primer for the amplification of *minD4* (*hvo_0322*) down-stream flanking region with a NdeI restriction site |
| 6918 | GACATCTAGAGTTGCGGCATCGCAGACGAG | Reverse primer for the amplification of *minD4* (*hvo_0322*) down-stream flanking region with a XbaI restriction site |
| 6919 | CGATGGTACCGTGGAGGTTGCGACGTTCTG | Forward primer for the amplification of *minD2* (*hvo_0595*) up-stream flanking region with a KpnI restriction site |
| 6920 | GTTCCATATGCCCACGTGCCCGGCGATTTC | Reverse primer for the amplification of *minD2* (*hvo_0595*) up-stream flanking region with a NdeI restriction site |
| 6921 | GTACCATATGAGAATCCCGCGGGGCGAACTC | Forward primer for the amplification of *minD2* (*hvo_0595*) down-stream flanking region with a NdeI restriction site |
| 6922 | GGACTCTAGAACCGTCGAGAAGCGTGTCAG | Reverse primer for the amplification of *minD2* (*hvo_0595*) up-stream flanking region with a XbaI restriction site |

| | | |
|---|---|---|
| 6923 | GTCAGGTACCGACCAGATCATCCCGACTTC | Forward primer for the amplification of *minD3* (*hvo_1634*) up-stream flanking region with a KpnI restriction site |
| 6924 | GACTCATATGCGGGGTTGGCCCCGGCTTGG | Reverse primer for the amplification of *minD3* (*hvo_1634*) up-stream flanking region with a NdeI restriction site |
| 6925 | CTAGCATATGTAATATGAGCGACCCCCTTAC | Forward primer for the amplification of *minD3* (*hvo_1634*) down-stream flanking region with a NdeI restriction site |
| 6926 | CCATTCTAGACACGTCGGCCATGTGTTCTG | Reverse primer for the amplification of *minD3* (*hvo_1634*) down-stream flanking region with a XbaI restriction site |
| 6583 | TGATTCTCGCGGTCGTCTCC | Forward primer to amplify a DNA probe for *minD1* |
| 6584 | AGGTTCGCCATTCCGAGGTC | Reverse primer to amplify a DNA probe for *minD1* |
| 6587 | TCTCGGTCCGAGTCGACAAC | Forward sequencing primer for Δ*minD1* |
| 6588 | TCTCGGAGTACCGCATCGAC | Reverse sequencing primer for Δ*minD1* |
| 6593 | GTTGTCGGATGTGTCATCTC | Forward primer to amplify a DNA probe for *minD3* |
| 6594 | AGTACTGGCGACGACTCCTC | Reverse primer to amplify a DNA probe for *minD3* |
| 6595 | TTACCAGCCTGTCGCTCCTC | Forward sequencing primer for Δ*minD3* |
| 6596 | GAGCCGTCTATCTTCGTGTC | Reverse sequencing primer for Δ*minD3* |
| 8029 | AATCGGAGGCCGAACCGAAC | Forward primer to amplify a DNA probe for *minD4* |
| 8030 | GTCGGTGTCCGAATCGACTG | Reverse primer to amplify a DNA probe for *minD4* |
| 8081 | CTAGGATCCTCAGACGAGCGAGGTGGTGTAGA | Reverse primer for the amplification of truncated *hvo_0322* with a Stop codon and BamHI restriction site |
| 8088 | GACGTGTTGCTCCTCGATTC | Forward primer to amplify a DNA probe for *minD2* |
| 8089 | TTTCAGCCCGTCAGACAGAG | Reverse primer to amplify a DNA probe for *minD2* |
| 8090 | CGAAAGCGAACGATTGGATG | Forward sequencing primer for Δ*minD2* |
| 8091 | AAGACGACGTAGCCGGTGAG | Reverse sequencing primer for Δ*minD2* |

| | | |
|---|---|---|
| 8826 | CGGGGTTGGAGCCACGACGAGCAC | Forward primer for the generation of the WalkerA mutation in MinD4 (on plasmid 3924) |
| 8827 | TGCTCGTCGTGGCTCCAACCCC | Reverse primer for the generation of the WalkerA mutation in MinD4 (on plasmid 3924) |
| 8828 | GTCATCATCGCCACCGGTGCG | Forward primer for the generation of the WalkerB mutation in MinD4 (on plasmid 3924) |
| 9728 | GATGTCGTAGCTCGCTTC | Reverse primer for the generation of the WalkerB mutation in MinD4 (on plasmid 3924) |
| 8847 | GATATCGAATTCCTGCAGCCCGGGGGATCCGCGTCGACCGCGCCGGAGAAC | Forward primer for the amplification of 525 Bp up-stream part of *minD4* with a 30 Bp overhang that is homolog to linearized pTA131 |
| 8848 | CTCCAGTGAAAAGTTCTTCTCCTTTACTCATAATTCCAACGTCGAAGTCCACCC | Reverse primer for the amplification of 525 Bp up-stream part of *minD4* with a 30 Bp overhang that is homolog to the start of *gfp* |
| 8849 | CTGTCACGGGTGGACTTCGACGTTGGAATTATGAGTAAAGGAGAAGAACTTTTC | Forward primer for the amplification of a 1282 Bp *gfp-minD4* part with a 30 Bp overhang that is homolog to the *gfp* part of primer 8848 |
| 8850 | GGTGGCGGCCGCTCTAGAACTAGTGGATCCAGGACGACGCCGACCACGTC | Reverse primer for the amplification of a 1282 Bp *gfp-minD4* part with a 30 Bp overhang that is homolog to linearized pTA131 |
| 8854 | GGTACCGGAACGACTGAATC | Forward sequencing primer for Δ*minD4* |
| 8855 | CGCCGTACATCCAGTACGTC | Reverse sequencing primer for Δ*minD4* |
| 8871 | GATAGGTACCCGACCGAGCGCGTCCTCTTC | Forward primer for the amplification of *pilB3*(*hvo_1034*) up-stream flanking region with a KpnI restriction site |
| 8872 | GTAGGATCCCGTCACCCGGTCAGTTGGTC | Reverse primer for the amplification of *pilB3*(*hvo_1034*) up-stream flanking region with a BamHI restriction site |
| 8873 | GTAGGATCCGATGGTCGCACAGTACCTGTATC | Forward primer for the amplification of *pilB3*(*hvo_1034*) down-stream flanking region with a BamHI restriction site |
| 8874 | GTACTCTAGAAGAGCACGTCGGTGCCGAAG | Reverse primer for the amplification of *pilB3*(*hvo_1034*) down-stream flanking region with a XbaI restriction site |
| 8889 | AGACGGCGCTGGTCGAGGAG | Forward primer to amplify a DNA probe for *pilB3* |

| | | |
|---|---|---|
| 8890 | CGCGCCGGAGGTAGTACAAC | Reverse primer to amplify a DNA probe for *pilB3* |
| 8893 | GCCGACGAGAGCGACCTGAC | Forward sequencing primer for Δ*pilB3* |
| 8894 | CGCGTCGCCATCGTCTGGAG | Reverse sequencing primer for Δ*pilB3* |
| 9767 | CTAGCTAGCTCAGACGAGCGAGGTGGTG TAGA | Reverse primer to amplify truncated *minD4* with a NheI restriction site |
| SI48 | GTATGGATCCTTAGTCGTCGGCGTCCGG GTCGGCCTCAG | Reverse primer to amplify truncated *minD4* (ΔS536-G574) with a BamHI restriction site |
| SI49 | CTAGCCGCGATGAGAAGAAAAAGGGGTT CCTCAGTCGCTTCCTCGGCTAAG | Oligonucleotide complementary to SI50 oligonucleotide with a NheI restriction site |
| SI50 | GATCCTTAGCCGAGGAAGCGACTGAGGA ACCCCTTTTTCTTCTCATCGCGG | Oligonucleotide complementary to SI49 oligonucleotide with a BamHI restriction site |
| SI75 | TTAGGCTAGCACGCCGGTCCCGGTTCCG GG | Forward primer to amplify the C-terminal part of *minD4* (T233-G574) |
| SI76 | GTACGGATCCTTAGCCGAGGAAGCGACT GA | Reverse primer to amplify the C-terminal part of *minD4* (T233-G574) |
| SI77 | CTAGCTCACCCGAAGACGACGAGGAGGG CGTCTACACCACCTCGCTCGTCGAGGAG GTCGAGTCGTTCGACGACGACGAGCGCG ATGAGAAGAAAAAGGGGTTCCTCAGTCG CTTCCTCGGCTAAG | Oligonucleotide complementary to SI78 oligonucleotide with a NheI restriction site |
| SI78 | GATCCTTAGCCGAGGAAGCGACTGAGGA ACCCCTTTTTCTTCTCATCGCGCTCGTCGT CGTCGAACGACTCGACCTCCTCGACGAG CGAGGTGGTGTAGACGCCCTCCTCGTCG TCTTCGGGTGAG | Oligonucleotide complementary to SI77 oligonucleotide with a BamHI restriction site |
| SI113 | TACGGGATCCTTATCCGGTGAGCGCCTC GGCGA | Reverse primer to amplify truncated *minD4* (ΔT233-G574) with a BamHI restriction site |
